# Informing epidemic (research) responses in a timely fashion by knowledge management - a Zika virus use case

**DOI:** 10.1101/2020.04.17.044743

**Authors:** Angela Bauch, Johann Pellet, Tina Schleicher, Xiao Yu, Andrea Gelemanović, Cosimo Cristella, Pieter L. Fraaij, Ozren Polasek, Charles Auffray, Dieter Maier, Marion Koopmans, Menno D. de Jong

## Abstract

The response of pathophysiological research to emerging epidemics often occurs after the epidemic and, as a consequence, has little to no impact on improving patient outcomes or on developing high-quality evidence to inform clinical management strategies during the epidemic. Rapid and informed guidance of epidemic (research) responses to severe infectious disease outbreaks requires quick compilation and integration of existing pathophysiological knowledge. As a case study we chose the Zika virus (ZIKV) outbreak that started in 2015 to develop a proof-of-concept knowledge repository. To extract data from available sources and build a computationally tractable and comprehensive molecular interaction map we applied generic knowledge management software for literature mining, expert knowledge curation, data integration, reporting and visualisation. A multi-disciplinary team of experts, including clinicians, virologists, bioinformaticians and knowledge management specialists, followed a pre-defined workflow for rapid integration and evaluation of available evidence. While conventional approaches usually require months to comb through the existing literature, the initial ZIKV KnowledgeBase (ZIKA KB) was completed within a few weeks. Recently we updated the ZIKA KB with additional curated data from the large amount of literature published since 2016 and made it publicly available through a web interface together with a step-by-step guide to ensure reproducibility of the described use case (S4). In addition, a detailed online user manual is provided to enable the ZIKV research community to generate hypotheses, share knowledge, identify knowledge gaps, and interactively explore and interpret data (S5). A workflow for rapid response during outbreaks was generated, validated and refined and is also made available. The process described here can be used for timely structuring of pathophysiological knowledge for future threats. The resulting structured biological knowledge is a helpful tool for computational data analysis and generation of predictive models and opens new avenues for infectious disease research.

**Availability:** www.zikaknowledgebase.eu

**Funding:** European Commission’s Seventh Framework Research Programme project PREPARE (FP7-Health n°602525) and ZIKALLIANCE (MK, H2020; No 734548).

**Author summary:** During the recent ZIKV outbreak there was little information about the interactions between Zika virus and the host, however, the massive research response lead to a steep increase in the number of relevant publications within a very short period of time. At the time, there was no structured and comprehensive database available for integrated molecular and physiological data and knowledge about ZIKV infection. Researchers had to manually review the literature (amounting to over 5000 articles on ZIKV during our last update of the ZIKA KB in September 2018) to extract information about host–pathogen interaction and affected molecular, cellular and organ pathways. We explored the use of automated literature analysis and a defined cooperative effort between experts from various scientific, biomedical and information-technology domains to rapidly compile existing pathophysiological knowledge as a potential tool to support investigations during an emergency. This tool is contrasted with conventional approaches that would take months to comb through the massive amount of existing literature. In addition to providing background information for research, scientific publications can be processed to transform textual information into complex networks, which can be integrated with existing knowledge resources to suggest novel hypotheses that potentially contribute to innovative infectious disease research approaches. This study shows that the knowledge extraction and mapping process required to inform clinical and research responses to an emerging epidemic can be efficiently and effectively executed with a dedicated and trained group of experts, a validated process and the necessary tools. Our results further provide an overview of ZIKV biology, allow prediction of drug efficacy and indentify specific host factors and signalling pathways affected by ZIKV.

## Introduction

The response to a (re-)emerging infectious disease (ID) epidemic requires a rapid compilation of existing pathophysiological knowledge to inform research priorities guiding basic and clinical research. Gaps in understanding of the underlying mechanisms make it difficult to design effective disease-modifying therapies. Hence, during an emerging ID outbreak, the available information at the time of its emergence and the subsequent rapid accumulation of scientific knowledge from various sources needs to be captured and analysed in a timely and comprehensive fashion. Responding to an ID outbreak therefore would benefit from the use of a knowledge repository that organizes the disease-related knowledge into pathway, molecular interaction and disease maps. Such maps are a relatively new concept that have been used in neurodegenerative and heart diseases (1,2), but which have had limited application in the field of ID thus far (3–5).

Molecular interaction and disease maps are dynamic computer-based knowledge repositories developed to integrate data and information across information sources, in a manner that is customized to the research domain of interest. Data types include interactions between molecular components, such as genes, pathogens, compounds and diseases.

The <u>P</u>latform fo<u>R</u> <u>E</u>uropean <u>P</u>reparedness <u>A</u>gainst (<u>R</u>e-)emerging <u>E</u>pidemics (PREPARE) is an EU-funded research consortium and clinical research network with the aim to rapidly respond to severe ID outbreaks, generating real-time evidence to inform optimized clinical management of patients and public health response. The 2015 ZIKV outbreak was considered as a test case in the context of the PREPARE network, as the pathogenesis of neurologic or immune disease induced by ZIKV is not fully understood. ZIKV is a flavivirus belonging to the *Flaviviridae* family and had only marginally been researched prior to the 2015 epidemic was minimal (6–8). Outbreaks of ZIKV disease have been recorded in Africa, the Americas, Asia and the Pacific. Acute ZIKV infections are mostly asymptomatic or associated with mild and self-limiting symptoms of fever, rash, conjunctivitis, headache or joint pain (9,10). However, the unexpected association of ZIKV infection with pregnancy and the subsequent severe neurodevelopmental problems in offspring and with the occurrence of neurological illnesses such as Guillain-Barre syndrome (GBS) or meningoencephalitis in acutely infected patients, led to widespread global concerns and a Public Health Emergency of International Concern (PHEIC) declaration by World Health Organisation (WHO) in 2016 (7).

We used the ZIKV virus outbreak as a case study to develop and test the steps, tasks, protocols and tools necessary to rapidly gather and integrate existing and emerging knowledge and to inform research priorities (Fig 1). Based on the available data and information we aimed to obtain a general overview of pathophysiological knowledge on ZIKV infection and its associated clinical manifestations described in the public domain. Other neurotropic flaviviruses, such as Dengue virus (DENV), West Nile virus (WNV), Japanese Encephalitis virus (JEV) and Tick-borne Encephalitis virus (TBEV) also cause nervous system infections, in particular encephalitis, but no association with neurodevelopmental disorders or GBS have been reported (11). To see whether including these viruses would shed additional light on ZIKV pathogenesis we compared available ZIKV information to other neurotropic flaviviruses in terms of neurovirulence and disease severity.

**Fig 1.**
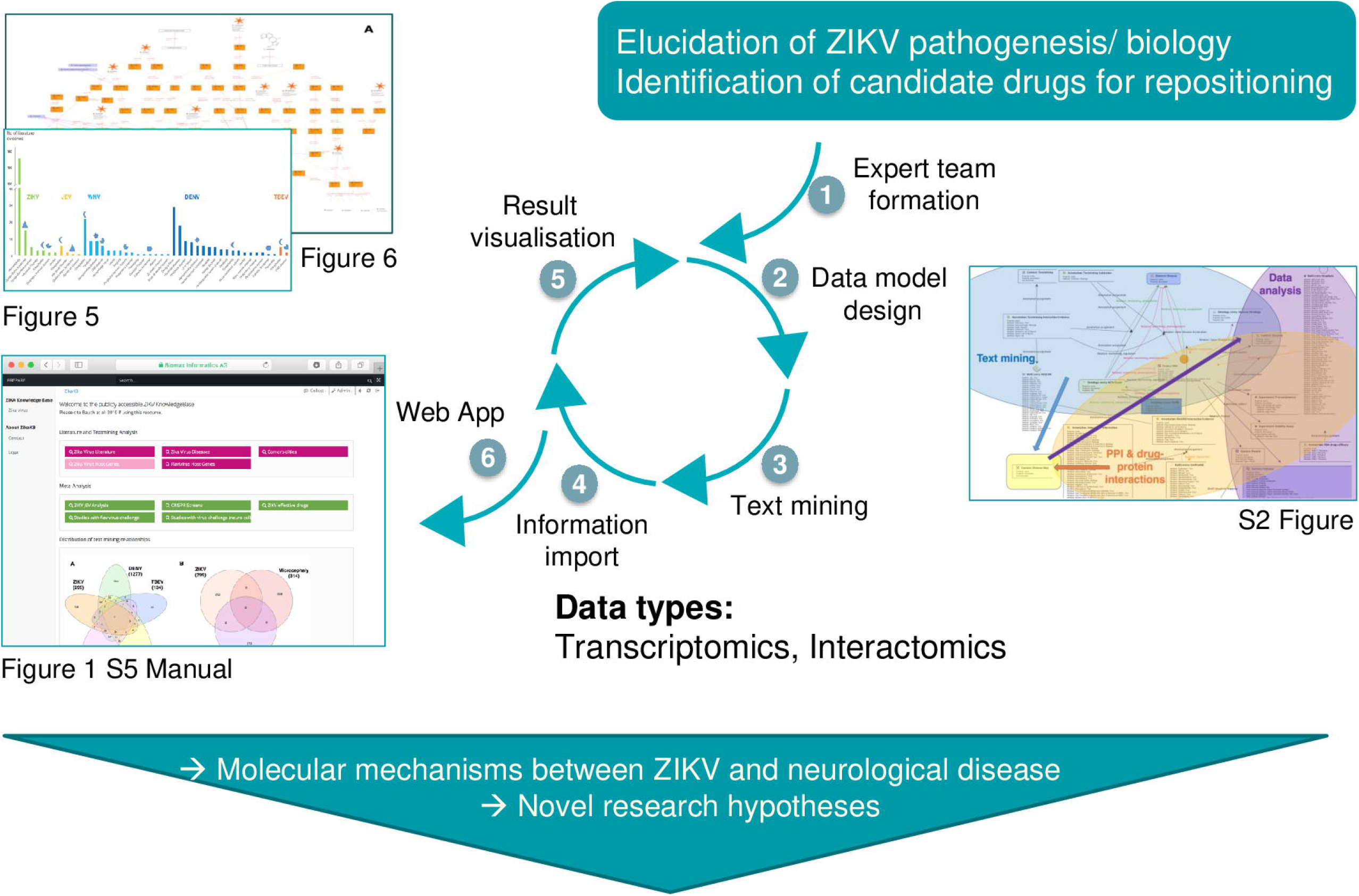
ZIKV KnowledgeBase generation process — Overview.

Based on the research objectives and knowledge provided by clinical/virology domain experts a six-step process was applied. In the first step a multidisciplinary expert team is assembled, in step 2, a semantic representation (“data model”) was designed by the knowledge management experts. This model includes details about the data sources for integration, how to transfer data into the system and how to report, visualize and export results, as well as the definition of the semantic context for objects, such as “gene”, “cell type” and “strain”. In a third step, a natural language processing algorithm was applied to the integrated PubMed literature source. In step 4 the relevant data, including literature mining results, was imported into the system and semantically mapped to the data model. In step 5, queries, views and reports were formulated. In the last step a web-browser based user interface was implemented to enable clinical/virology experts to review, validate and refine the integrated information.

## Methods

### Rapid response protocol

Many procedures have been published to collect knowledge from literature and experts, including systematic literature reviews (12), clinical guideline consensus building (13) and literature mining (14). Based on these approaches we developed a dedicated six step protocol with a focus on rapid assembly of existing knowledge (see Fig 1 and S1 Fig):

#### 1. Team organisation and process management

A multidisciplinary team of clinicians, virologists, bioinformaticians and knowledge management specialists was formed to collaboratively extract existing ZIKV related knowledge from the literature and from public databases, integrate the available information into a consistent summary and further connect integrated data to molecular and pharmaceutical information. To enable an efficient and consolidated initial result, tasks were distributed between individuals and results were discussed and integrated in weekly online conferences. The detailed protocol for knowledge base generation was developed in this initial exercise and is presented in the results section.

#### 2. Knowledge Management

For data organization, integration and development of molecular interaction and disease maps, a dedicated knowledge management tool is required. In this case, one of the PREPARE partners contributed the BioXM™ Knowledge Management Environment, a generic platform for dynamic modelling, visualization and analysis of biological and biomedical networks (15). For knowledge representation we applied a semantic network approach as described previously (16,17). Briefly, the abstract concepts necessary to capture and represent essential ideas and physical objects relevant to the domain of knowledge were defined. Based on input from clinical and virology experts, the concepts required to represent existing pathophysiological knowledge of infectious diseases were modelled with objects, such as “genes”, “strains”, “is expressed in” or “interacts with”. For the ZIKV KB, we focused on concepts required to represent text-mining results and information from structured databases of protein–protein (PPI) and drug–protein interactions, namely genes, diseases, pathogens and drugs. Relationships between pathogens, genes, drugs and compounds extracted by text-mining were represented by three types of relations: up-regulation, down-regulation and regulation (for further details see S2 Fig). Where possible, each concept was referenced to unique entries from reference databases or ontologies such as ChEBI (18) for chemicals and ICD10 (19) for diseases. The defined semantic concepts become directly available in a natural-language-like query and reporting language. This language can be used to address specific questions and to summarise and visualise available knowledge. For example, the query “is a *drug* which *interacts with* is a *protein* from *organism human* which *is expressed by* a *gene* which is *associated with* a *organism Zika*” retrieves the number of drugs that interact with a protein of interest and generates a visualisation which applies a color coding to genes that indicates to the number of associated drugs.

#### 3. Text mining

The integrated text mining tool uses syntactic text parsing and dictionary-based named-entity recognition to extract semantically typed associations (such as “inhibits”, “activates”) between the defined semantic concepts (such as “gene”, “strain”) (20). The initial task creates a defined text corpus, which includes uploaded relevant full text articles if applicable. In principle the textual materials for mining can be derived from PubMed abstracts, text from the WHO or other news feeds or any document in the portable document format (PDF), Mircorsoft Word or American Standard Code for Information Interchange (ASCII) formats. For the case study described here we used all ZIKV PubMed abstracts and publicly openly available full text articles. From these sources, relationships between genes, diseases, pathogens and drugs were extracted. The extracted associations consist of a subject, an object and the linking predicate and are enriched by their supportive evidence and additional metadata. For example, one such relationship is “Zika virus (subject) causes (predicate) microcephaly (object)” (Fig 2). Genes, diseases, pathogens and drugs, can be used as subjects and as objects and the sum of all extracted associations form an initial knowledge network. Genes, diseases, pathogens and drugs were defined by dictionaries that we curated from public sources as described below. Each dictionary consists of a well-defined set of ontologies (including synonyms) or reference databases tailored to the research question of interest. For instance, the disease dictionary consists of Disease ontology entries (21) and relevant branches of the NCI Thesaurus (22), the organism dictionary of NCBI taxonomy entries (23), the compound dictionary of ChEBI entries (18), as well as of KEGG (24) and NCI Thesaurus compounds. The gene dictionary is based on genes derived from human and flavivirus genomes. Predicates are derived from a set of verbs, which can be modified. These predicates describe mainly molecular interactions but can also indicate causal associations between proteins or compounds and diseases (for instance “activates”, “restricts”, “targets”). To optimise recall and specificity of the mining, we extended the dictionaries for viral names, acronyms and interaction predicates as well as defined a black-list of acronyms causing mostly false positives.

**Fig 2.**
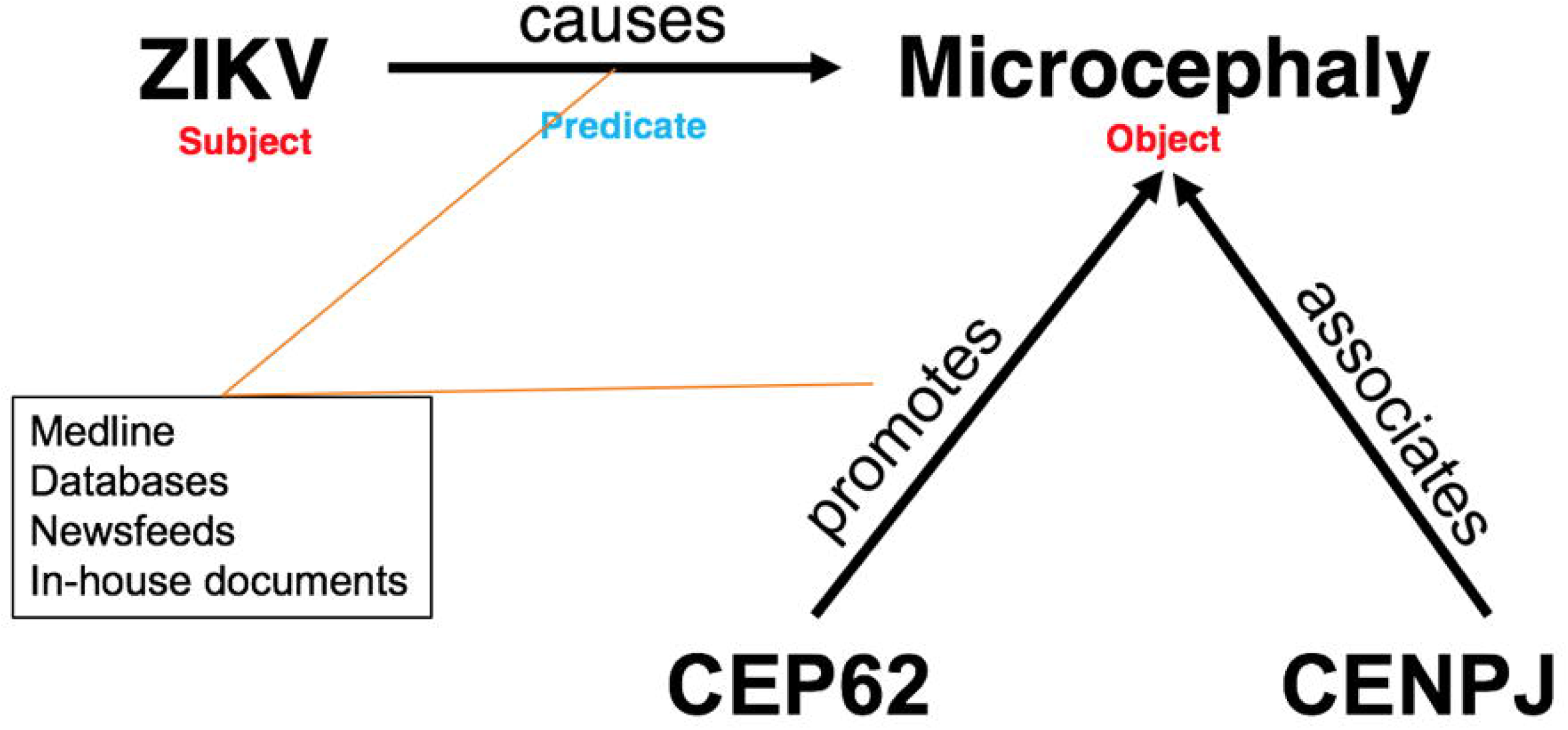
Predicated text mining relationship.

A text mining relationship consists of a subject (ZIKV), an object (microcephaly) and a linking predicate (causes). Subject and object are defined by dictionaries consisting of ontologies or reference databases, whereas predicates are derived from a fixed set of verbs with assertions from integrated sources, such as Medline. The term “microcephaly” is a reusable scientific concept that participates not just in one “Subject-Predicate-Object” construct detected, but in all such constructs detected that mention “ microcephaly”. Supplementary information is associated with the “microcephaly” object, including, for example, information from the Disease ontology, and other integrated resources, such as Gene-Disease-Association data (DisGeNET). This expandable set of relationships forms a large network of knowledge that enables new knowledge to be inferred by “reasoning” based on the logic encoded in those relationships.

Finally, the extracted relationships can be curated to manually optimise quality and information content. A curation user interface was implemented to enable the expert team to support or refute the automatically generated relationships. At least two independent researchers (the “4-eye review mode”) manually evaluated the evidence for every extracted relationship. In the case that the evaluations from the two researchers conflicted, the conflicts were either resolved during the weekly online conferences or were excluded, as our goal was to maximise specificity (correctness) rather than sensitivity (completeness) of the integrated information.

In addition, experts could expand the network with any relevant supporting evidence from other integrated sources, such as public or proprietary databases and experimental data.

#### 4. Semantic mapping of experimental results, public data sources and ontologies

Semantic mapping describes the process of identifying and linking concepts that are shared between two information sources. We integrated the databases listed in Table 1 using existing concepts such as genes, pathogens or diseases which were identified by ontological descriptors. Semantically identical objects are mapped to descriptive data from literature and databases to allow informed and efficient querying of the overall collected information (e.g. “Dengue disease” is mapped to the following synonyms: “Breakbone fever”, “Dengue disorder”, “Dengue fever” and “Dengue”) (25). To this end, mapping scripts are created to resolve a given input data format and match the provided entity identifiers or ontology terms. Experimental data from key publications is mapped by the same approach. While these data are henceforth available for search and reporting they are not yet displayed as part of any specific molecular interaction and disease map.

**Table 1.**
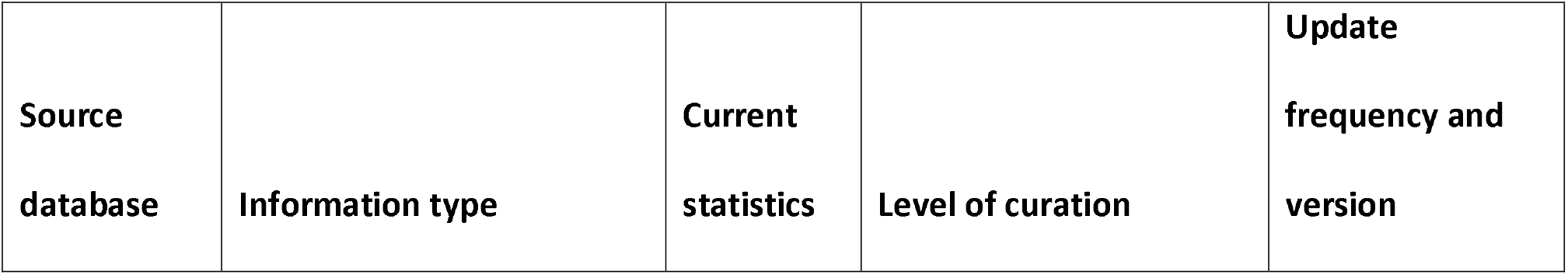
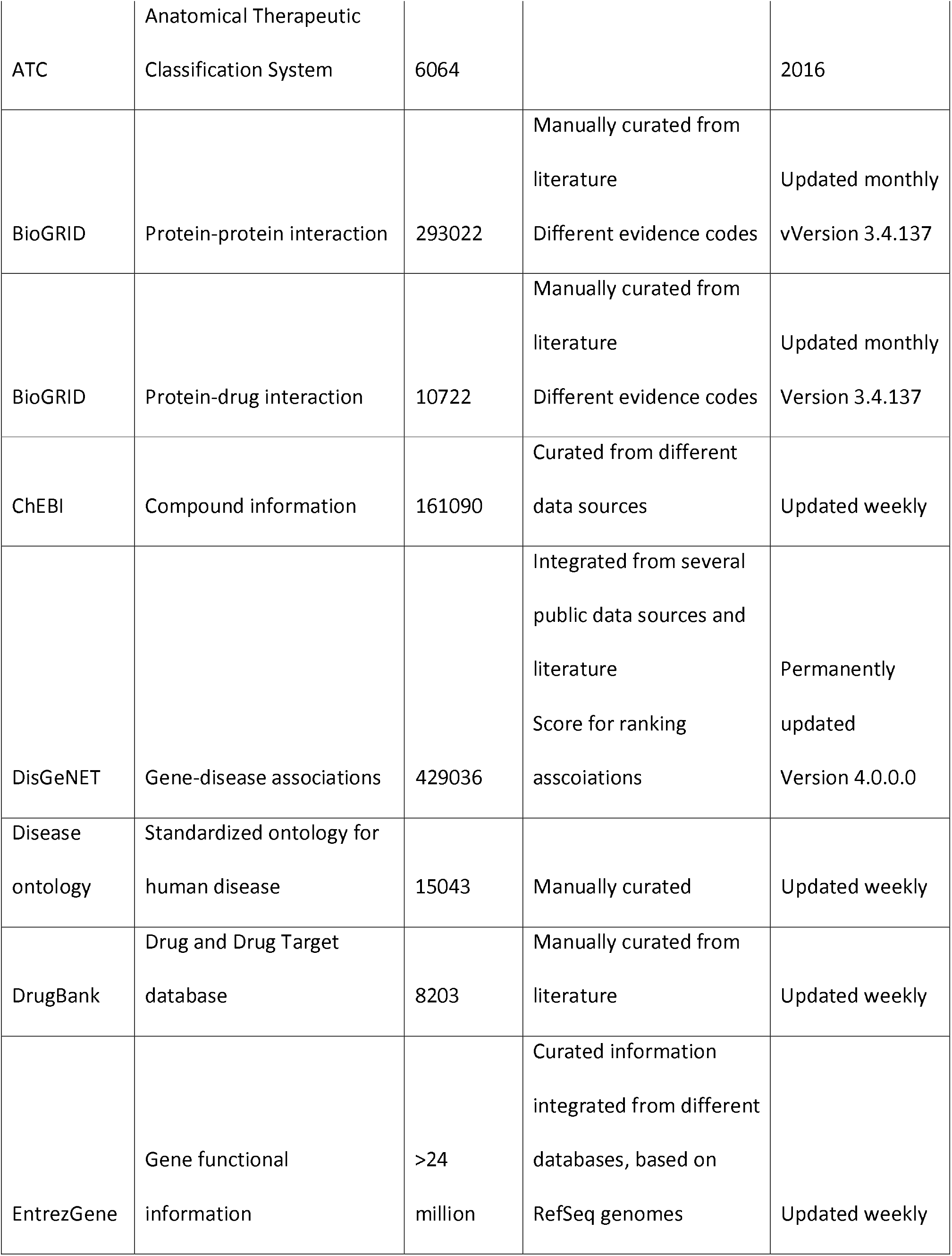
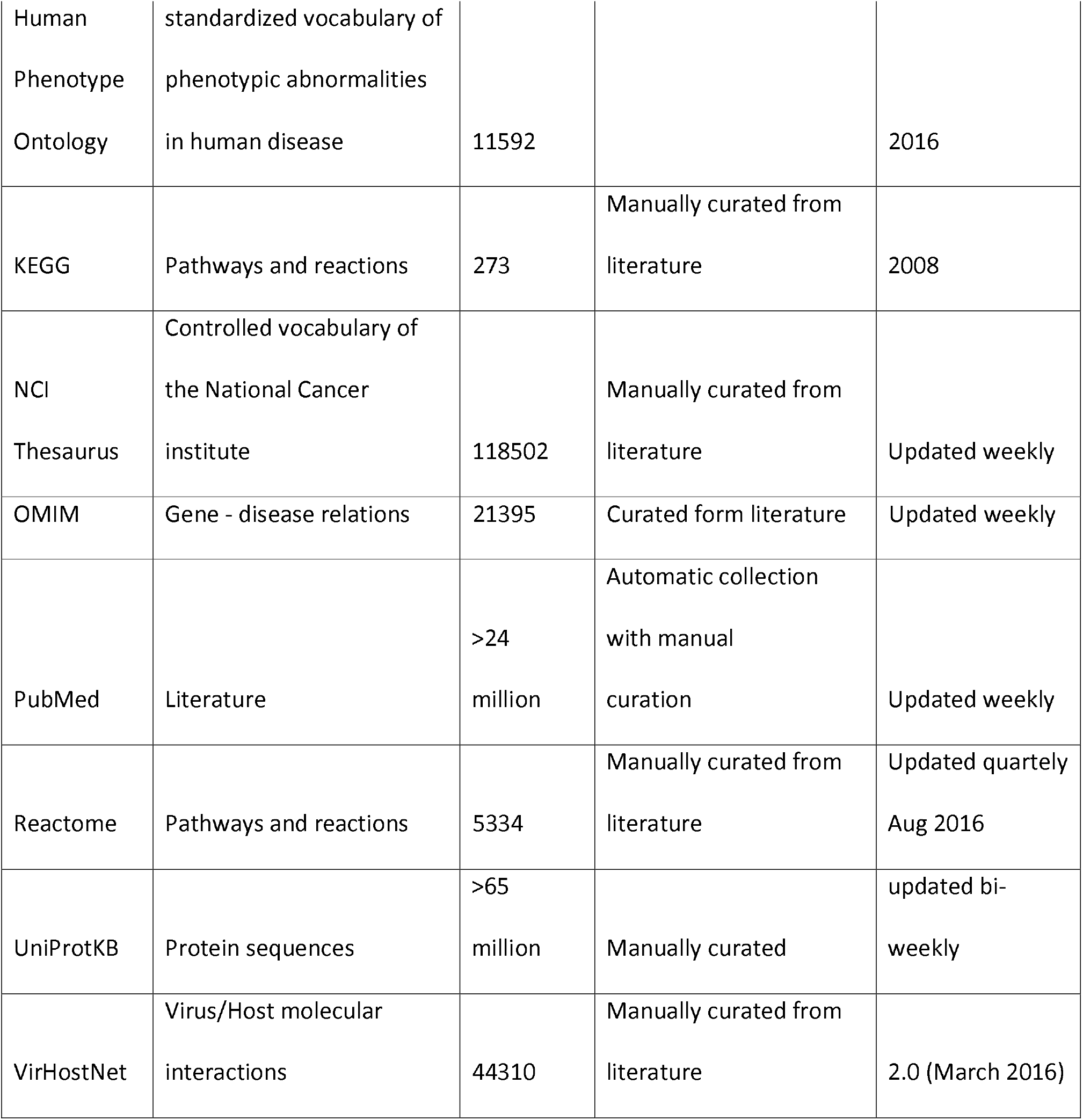
External data sources integrated into ZIKV KnowledgeBase.

| Source<br>database | Information type | Current<br>statistics | Level of curation | Update<br>frequency and<br>version |
| --- | --- | --- | --- | --- |
| ATC | Anatomical Therapeutic Classification System | 6064 |  | 2016 |
| BioGRID | Protein-protein interaction | 293022 | Manually curated from literature<br>Different evidence codes | Updated monthly<br>vVersion 3.4.137 |
| BioGRID | Protein-drug interaction | 10722 | Manually curated from literature<br>Different evidence codes | Updated monthly<br>Version 3.4.137 |
| ChEBI | Compound information | 161090 | Curated from different data sources | Updated weekly |
| DisGeNET | Gene-disease associations | 429036 | Integrated from several public data sources and literature<br>Score for ranking associations | Permanently updated<br>Version 4.0.0.0 |
| Disease ontology | Standardized ontology for human disease | 15043 | Manually curated | Updated weekly |
| DrugBank | Drug and Drug Target database | 8203 | Manually curated from literature | Updated weekly |
| EntrezGene | Gene functional information | >24 million | Curated information integrated from different databases, based on RefSeq genomes | Updated weekly |
| Human<br>Phenotype<br>Ontology | standardized vocabulary of<br>phenotypic abnormalities<br>in human disease | 11592 |  | 2016 |
| KEGG | Pathways and reactions | 273 | Manually curated from<br>literature | 2008 |
| NCI<br>Thesaurus | Controlled vocabulary of<br>the National Cancer<br>institute | 118502 | Manually curated from<br>literature | Updated weekly |
| OMIM | Gene - disease relations | 21395 | Curated form literature | Updated weekly |
| PubMed | Literature | >24<br>million | Automatic collection<br>with manual<br>curation | Updated weekly |
| Reactome | Pathways and reactions | 5334 | Manually curated from<br>literature | Updated quartely<br>Aug 2016 |
| UniProtKB | Protein sequences | >65<br>million | Manually curated | updated bi-<br>weekly |
| VirHostNet | Virus/Host molecular<br>interactions | 44310 | Manually curated from<br>literature | 2.0 (March 2016) |

#### 5. Querying and visualization of integrated information in tables, networks and disease maps

To help the expert team establish a specific molecular interaction and disease map we defined a number of queries to explore the collective knowledge. These queries were used, for example, to find diseases and genes associated with a virus of interest to find diseases associated with genes prioritized according to experimental evidence.

Based on these queries we developed a streamlined, wizard-based user interface to create disease maps by selecting the relevant relationships from the curated text mining from query results (S3 Fig). This basic network was further extended with interaction data (e.g. PPI & protein-drug interaction) by applying a network search algorithm based on genes extracted from text mining relationships. Finally, we defined queries to overlay additional information, such as literature evidence, experimental data, drug targets or host factors to obtain different perspectives of the same underlying molecular interaction or disease map.

#### 6. Deployment of an open access, web-based user interface

To make the results of our internal test case generally available and to support ZIKV research, we provide and maintain a regularly updated ZIKA KB at the following URL www.zikaknowledgebase.eu. As we continue to extend this resource user registration for access will be implemented to ensure the knowledge base is used for research only.

## Results

### Semantic representation of ZIKV infection

The data model implemented to provide a semantic representation of ZIKV infection is described in detail in supplemental Figure S2. Briefly, the model focuses on genes, diseases, pathogens and drugs, and distinguishes between associations derived from literature mining and those provided by experimental data such as PPIs.

### Text mining results

We searched PubMed with the terms “Zika virus”, “Dengue”, “West Nile virus”, “Japanese encephalitis virus”, “Tick-borne encephalitis virus”, “Microcephaly” and “Guillain Barre Syndrome” initially in December 2016 and most recently in September 2018. The recent search resulted in 4927 hits for “Zika virus” and 19974, 7700, 5918, 5213, 14248 and 8615 hits for the other search terms, respectively. During the analysed time frame, literature on ZIKV increased substantially from 1414 in 2016 to the current 4927 hits (250%), whereas for all other terms, the increase in publications was closer to 10%. Accordingly, the recent search identified additional disease phenotypes, including carditis and skin diseases, that were reported to be associated with ZIKV that were not present in the previous search. An additional set of 236 open access full text articles about ZIKV were included. A natural language processing algorithm was applied to these sets of documents to efficiently extract the fast growing information in the biomedical literature. The text mining extracted a total of 11916 relationships, which were manually evaluated to 2982 verified relationships (Table 2). The distribution of the curated relationships is depicted in Fig 3, indicating that the largest overlap was for ZIKV and DENV and for DENV and WNV. The curated set of relationships was used for further analyses, including generation of molecular interaction and disease maps and querying for virus-associated genes or diseases.

**Table 2.**
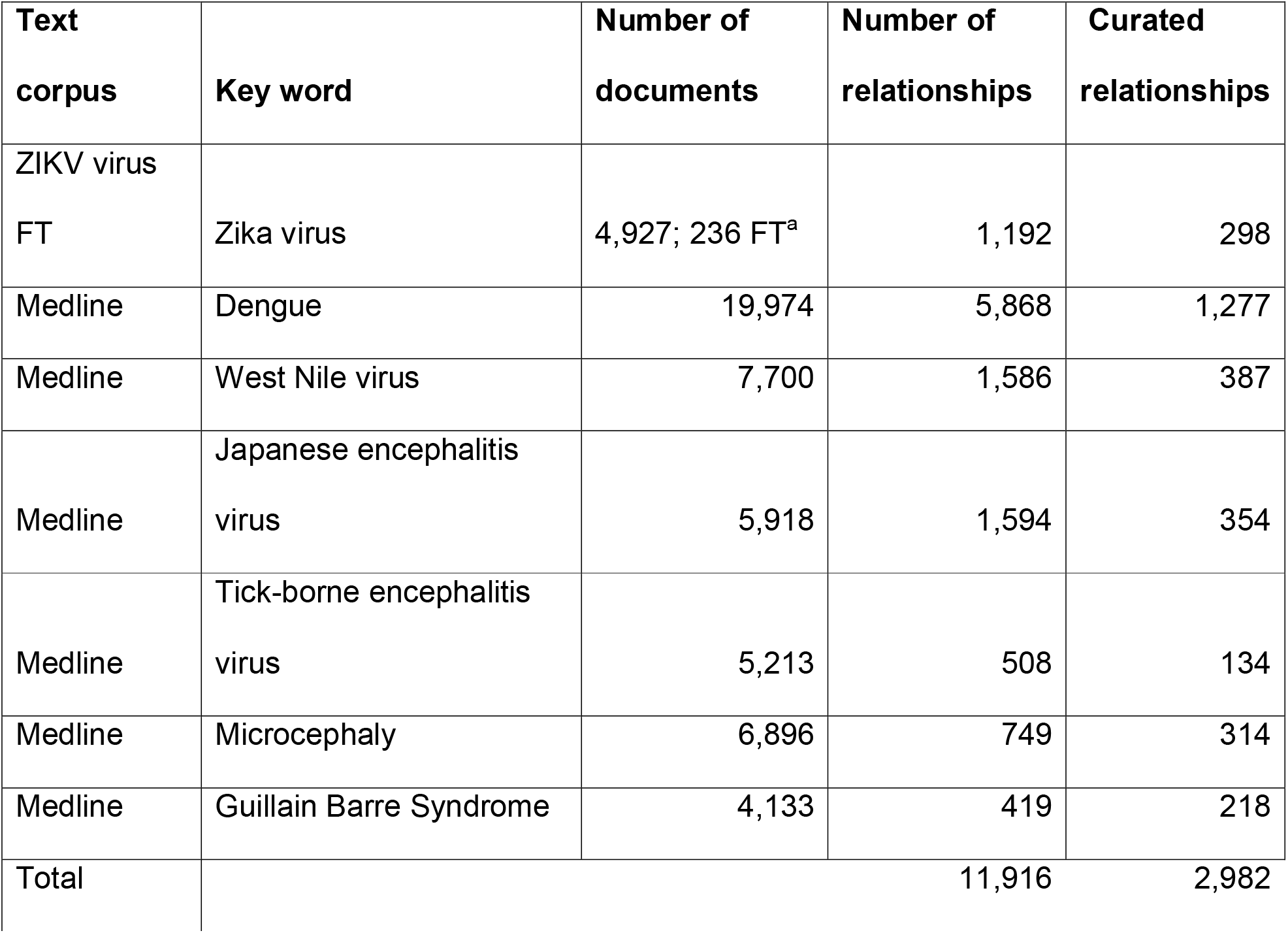
Text mining analyses. ^a^FT: full text.

**Fig 3.**
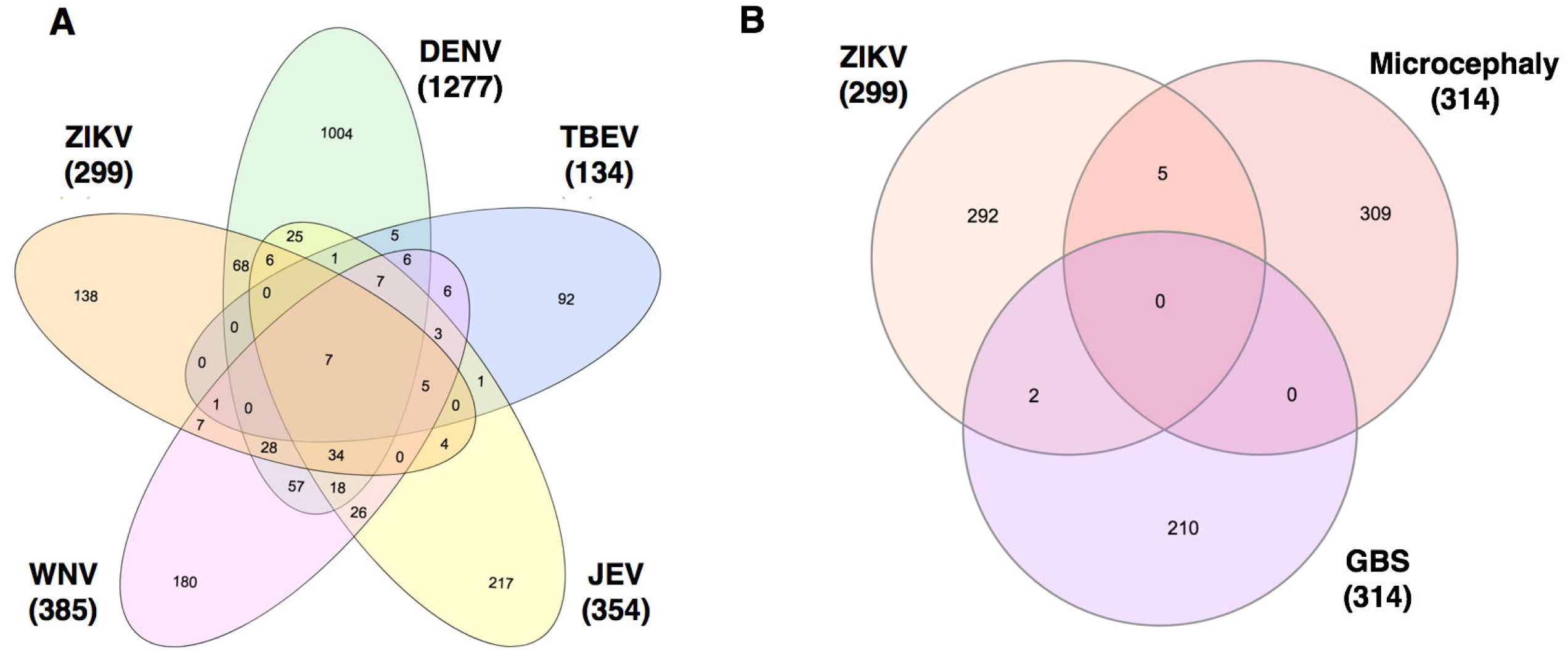
Distribution of text mining relationships. Numbers represent the sum of different types of relationships, such as gene-pathogen, gene-disease, gene-gene, gene-compound and pathogen-disease relations, found for each virus in total as well as in overlap with other viruses (A), or ZIKV in overlap with text mining analyses of Microcephaly and Guillain-Barre Syndrome (B).

### Integrated data

Overall, the ZIKA KB contains a network of 337332 human and host-pathogen PPI integrated from BioGRID and VirHostNet, as well as 18905 protein-drug interactions integrated from BioGRID and DrugBank, and 450431 gene-disease associations from DisGeNET (Table 1). Recently, a variety of ZIKV- and other flavivirus-related large-scale data sets, including microarray gene expression (26,27), RNAseq (28) as well as CRISPR/Cas data (29), have become publicly available and were integrated to identify host factors that are affected during viral infection.

### Molecular interaction and disease maps

Curated text mining results were used to populate the initial ZIKV molecular interaction and disease map. In a second step the map was extended with interaction data (PPI & protein-drug interaction by applying, a network search to implement the breadth-first algorithm (30) which connected genes extracted from text mining relationships based on the overall network. This set of interaction data can be filtered and explored interactively. In a systems medicine approach, a multidisciplinary expert team systematically analysed literature, public databases and experimental resources to create a formal, structured model of molecular and cellular ZIKV–host interactions (“molecular interaction and disease map”)

### Publicly available ZIKA KB

After an assessment period of internal use, a web-browser based user interface was implemented to make the PREPARE ZIKA KB available to all ZIKV researchers. By openly sharing the collected data and information, the ZIKA KB allows researchers to generate hypotheses, identify knowledge gaps and interactively explore and interpret data. All data are currently in the public domain. Upon request, data submission can be modified to allow registered users to specify that submitted data should not be publicly available

### Use of the ZIKA KB

In the following we provide several example use cases. For instance, publicly available interaction data, such as the PPI and protein–drug interaction data can be used to visualize drug targets and host factors involved in ZIKV pathogenesis. Alternatively, users can filter for PPIs whose source or target is a drug or refine search results to include only proteins localized to a specific cellular compartment, such as the endoplasmic reticulum. The returned networks can be interrogated subsequently to identify host factors that are targeted by the virus and to search for drugs that interact with these host factors. and thus might contribute to drug repositioning for future treatment options for ZIKV infection. The maps can also be explored further by using integrated expression and knockout data.

To explore the integrated literature knowledge for relevance or obtain an overview of drug targets or identify critical genes within the network consisting of gene-disease-pathogen relationships, predefined perspectives were overlaid onto the default map. The association of ZIKV with microcephaly was reported most frequently across all ZIKV literature and this association is visualized by the thickness of the edges (Fig 4A). Known drug targets interacting directly with ZIKV or microcephaly were highlighted in green for potential intervention evaluation (Fig 4B). Genes playing a role in ZIKV infected human neural progenitor cells (hNPCs) were also highlighted for comparative analyses of complementary experimental analyses (Fig 4C).

**Fig 4.**
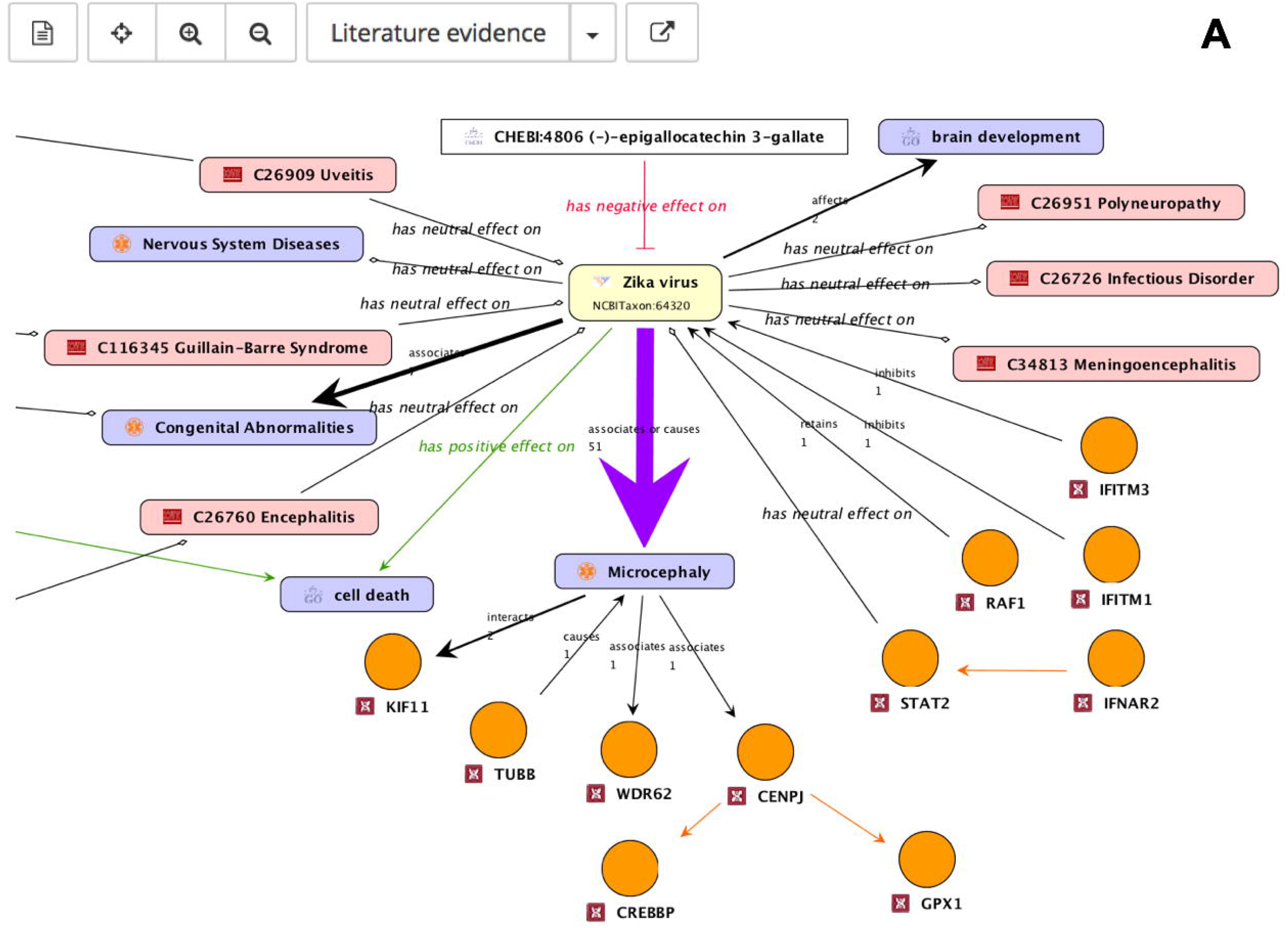
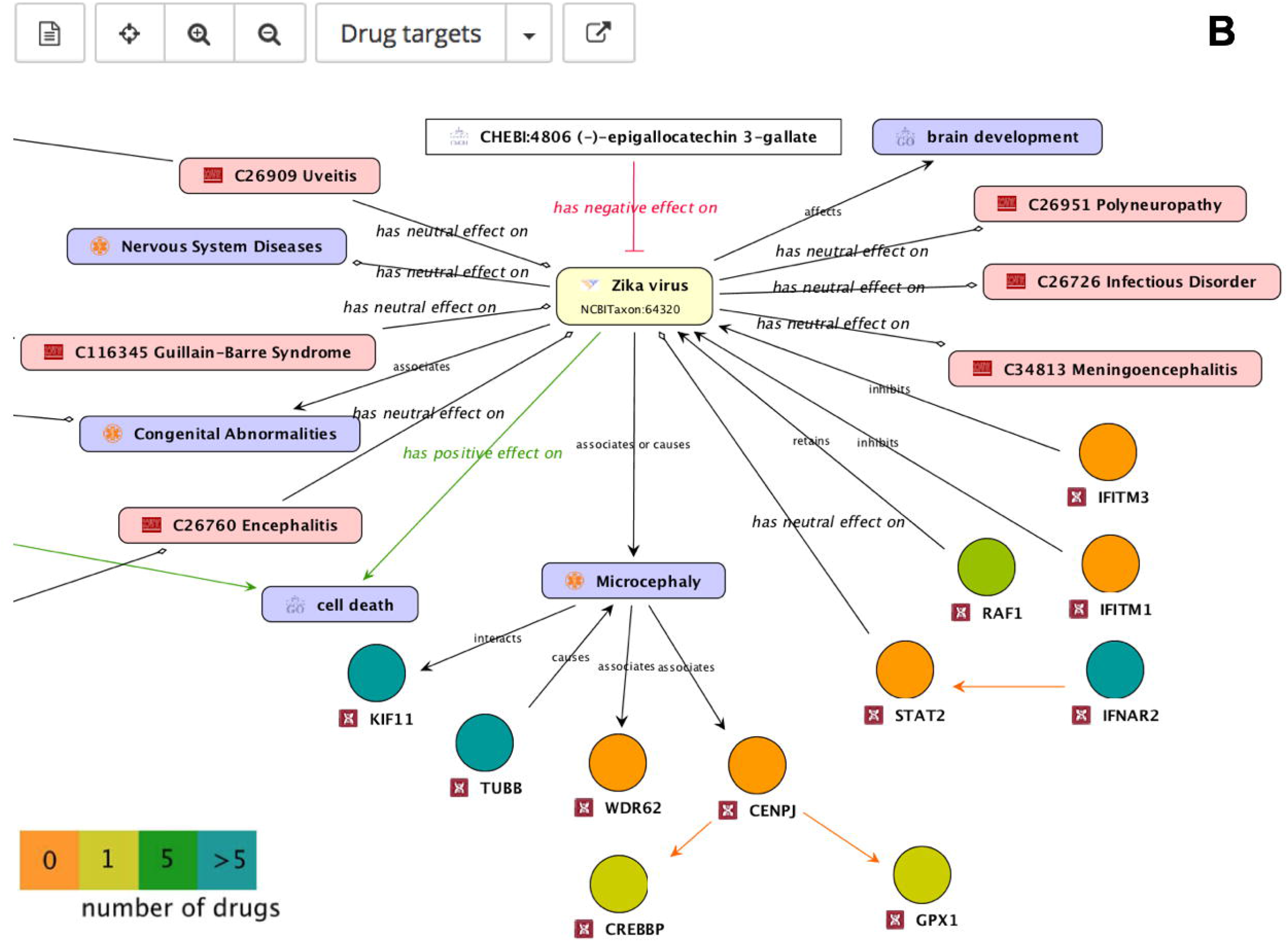
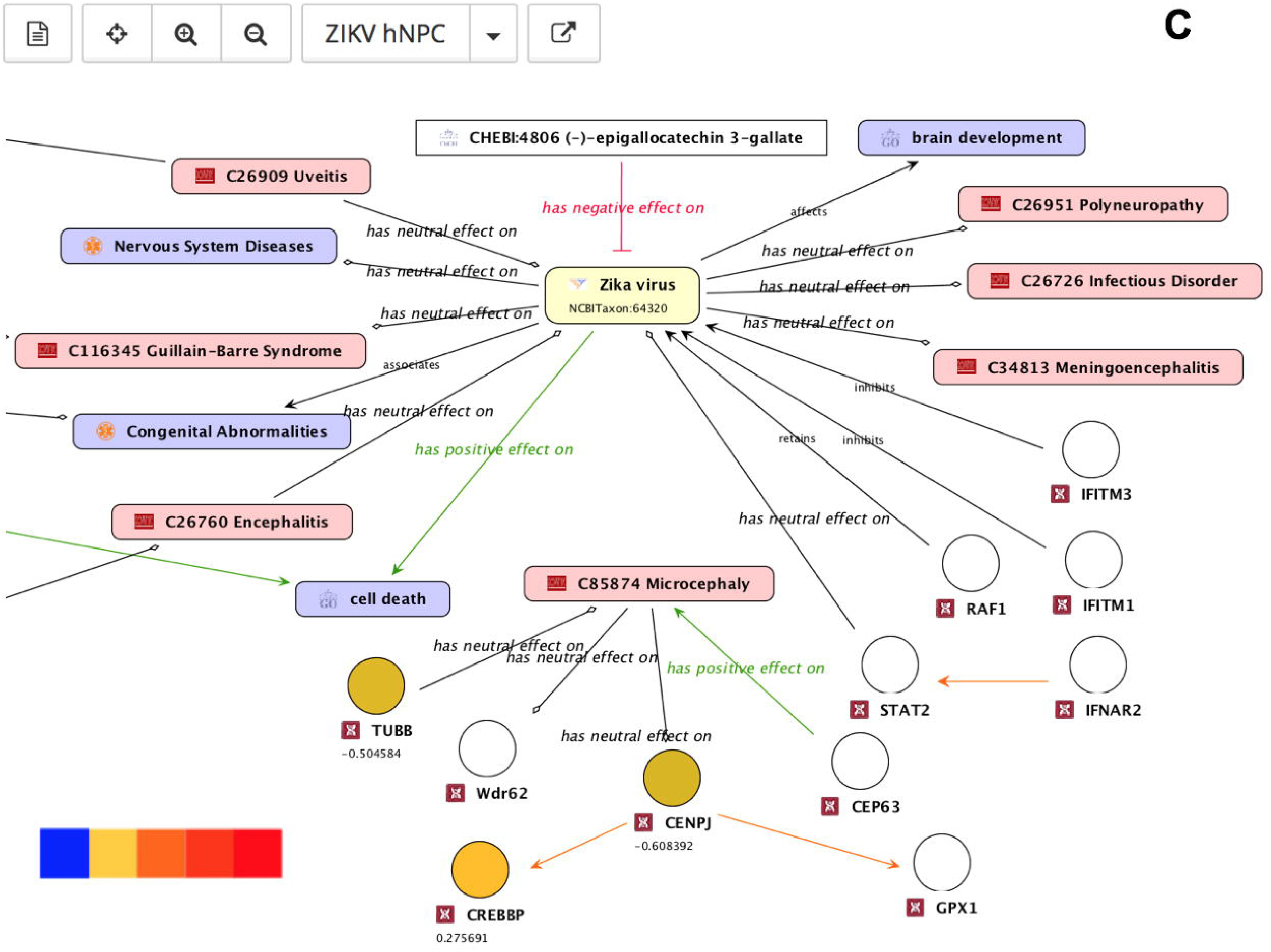
Different ZIKV molecular interaction and disease map perspectives. (A) The amount of literature evidence is depicted by relation strength. (B) Genes known to be drug targets are color coded: the gene is a target for one (green) or five or more (blue) drugs or no drugs (orange). (C) Genes (hNPCs challenged with ZIKV, Tang et al, 2016) are color coded to indicate up-(red) and down-(blue) regulation.

#### Diseases associated with flaviviruses

To further explore and validate the knowledge derived from the curated text mining analyses, diseases associated with a virus of interest were queried. The results are displayed in Fig 5. It was assumed that the disorders that were most frequently associated with a virus were the primary disorder for infection with the virus. Microcephaly and GBS, for instance, are the most frequently mentioned disorders associated with ZIKV infection. Dengue fever and hematopoietic system disorders (e.g. thrombocytopenia) are frequently listed for DENV, whereas encephalitis is the most frequently mentioned disease for WNV, JEV and TBEV. Each disease thus represents the corresponding primary manifestation of these viral infections. Encephalitis is equally frequently associated with ZIKV and DENV confirming that ZIKV is also an aetiological agent in encephalitis.

**Fig 5.**
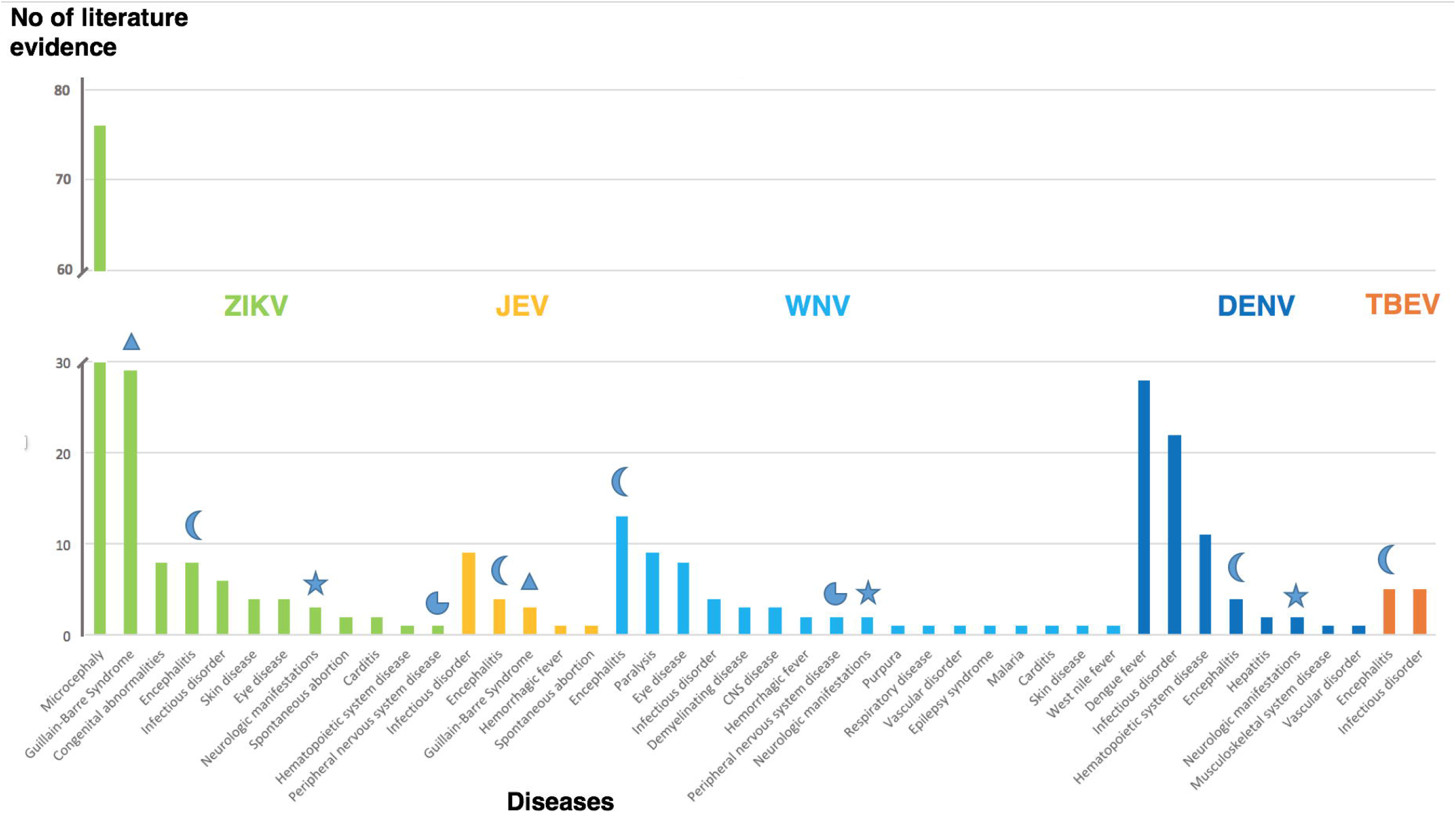
Diseases associated with flaviviruses. The amount of literature evidence (y-axis) for each of the diseases (x-axis) is grouped for the five flaviviruses. Symbols highlight neurological diseases associated with more than one virus. Triangle: GBS, moon: Encephalitis, pie-chart: Peripheral nervous system disease, star: Neurologic manifestations

#### Potential inhibitors of ZIKV infection

Currently there is no approved therapy to treat ZIKV infection. Barrows *et al* recently performed a screen of 774 FDA-approved drugs to identify agents that could potentially be repositioned as treatment options for ZIKV infection (31). Of these, 24 potential inhibitors of ZIKV infection were identified and validated in human neural stem cells and primary amnion cells. In addition to their potential use for treatment, these compounds provide a resource to study ZIKV pathogenesis and can contribute to insights into the biology of ZIKV. To this end, the ZIKV molecular interaction and disease map described in Figure 4 was extended and filtered to include these potential “*ZIKV effective drugs*” which were connected to genes associated to ZIKV through PPIs (Fig 6). After this extension ten of the identified *ZIKV effective drugs* were part of the new map which we then used to gain insight into potential drug mechanisms and ZIKV biology. One of the drugs, Bortezomib, is a known antiviral compound that inhibits replication of flaviviruses (32). Bortezomib is a proteasome inhibitor, suggesting that proteasome action is essential for ZIKV replication. This conclusion is in agreement with published CRISPR screen data (29) identifying genes associated with protein degradation required for ZIKV infectivity. Interestingly, four of the predicted *ZIKV effective drugs* (Mefloquine, Mebendazole, Sorafenib and Dactinomycin) are associated with genes which, through PPIs, are involved in ErbB signalling. *ErbB* is associated with the development of neurodegenerative diseases when inactivated (33). Four of these genes (*MYC*, *GSK3B*, *BRAF* and *MAP2K2*) are reported to be up-regulated in a published RNAseq analysis (28) performed in human embryonic cortical neural progenitor cells (Fig 7). These genes can serve as an entry point to be tested in specific assays designed to unravel molecular mechanisms between of ZIKV involvement in microcephaly.

**Fig 6.**
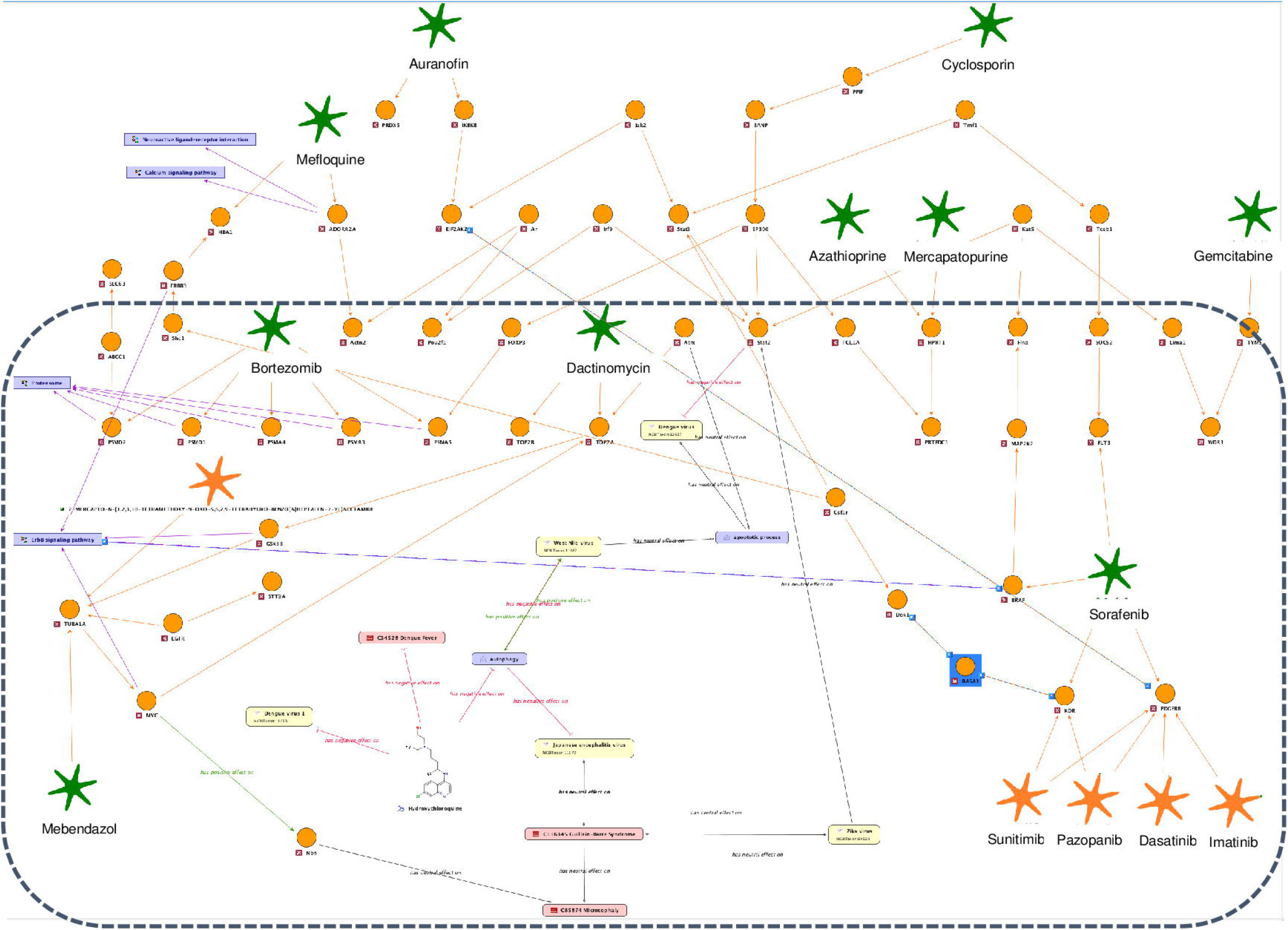
ZIKV molecular interaction and disease map extended for ZIKV effective drugs. The following are the symbols used in the map. Orange circles: genes; green stars: ZIKV effective drugs; yellow rectangles: flaviviruses; pink rectangles: diseases; violet rectangles: GO processes or KEGG signalling pathways. The following edge colors are used in the map. Black edges: relationships derived from text mining; orange: protein-drug or PPIs. The latter were obtained by applying a network search algorithm selecting the drug target of a ZIKV effective drug as start and STAT2 (contained in a direct relation with ZIKV) as the end point.

**Fig 7.**
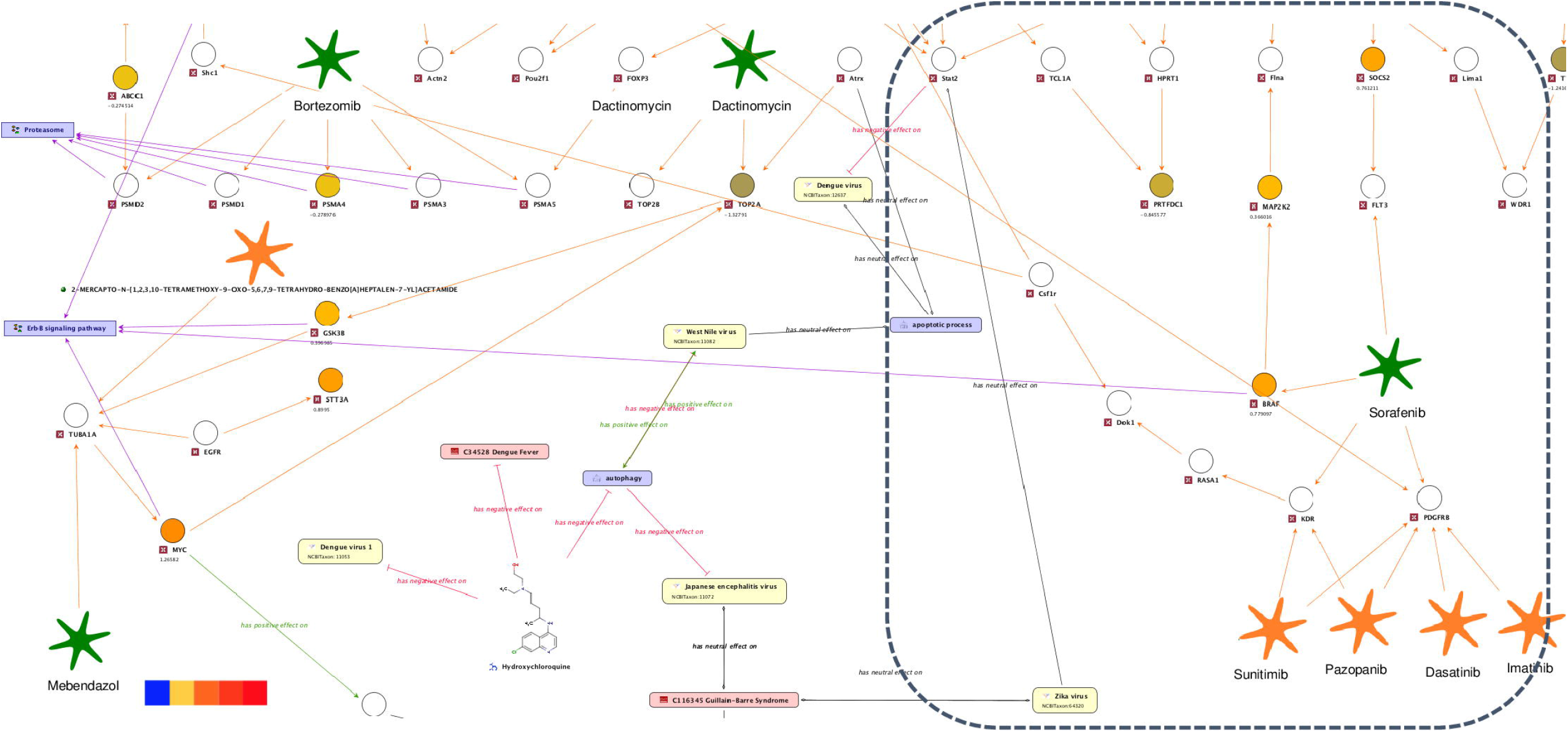
ZIKV molecular interaction and disease map extended for ZIKV effective drugs (enlarged perspective). Up- and down-regulated genes (hNPCs challenged with ZIKV, Tang et al, 2016) are highlighted by a color code ranging from red to blue from enlarged perspectives surrounded by dotted boxes in Figure 6.

Another predicted *ZIKV effective drug* is Sorafenib, a multi-target tyrosine kinase inhibitor. The ZIKV map was used to identify the effective target of Sorafenib:

1. 1. Sorafenib interacts with 4 target genes, *FLT3*, *BRAF*, *VEGFR* (also known as *KDR*) and *PDGFR*. The latter two genes are known to interact with additional drugs, such as Sunitinib, Pazopanib, Dasatinib and Imatinib. These additional drugs were among those which had no effect in the ZIKV infection assay.
2. In ZIKV infection expression data, none of the genes producing protein products that interact with VEGFR and PDGFR through known PPI (orange edges) with ZIKV are differentially expressed (Fig 8). In contrast, *BRAF* and *SOCS2*, a FLT3 interactor, were unregulated upon ZIKV infection. Based on the observations above drawn from the ZIKV map, we hypothesise that FLT3 or BRAF are the effective targets of Sorafenib in ZIKV infection, rather than VEGFR and PDGFR. This exemplifies how molecular interaction and disease maps can be used to provide further insight into ZIKV biology.

**Fig 8.**
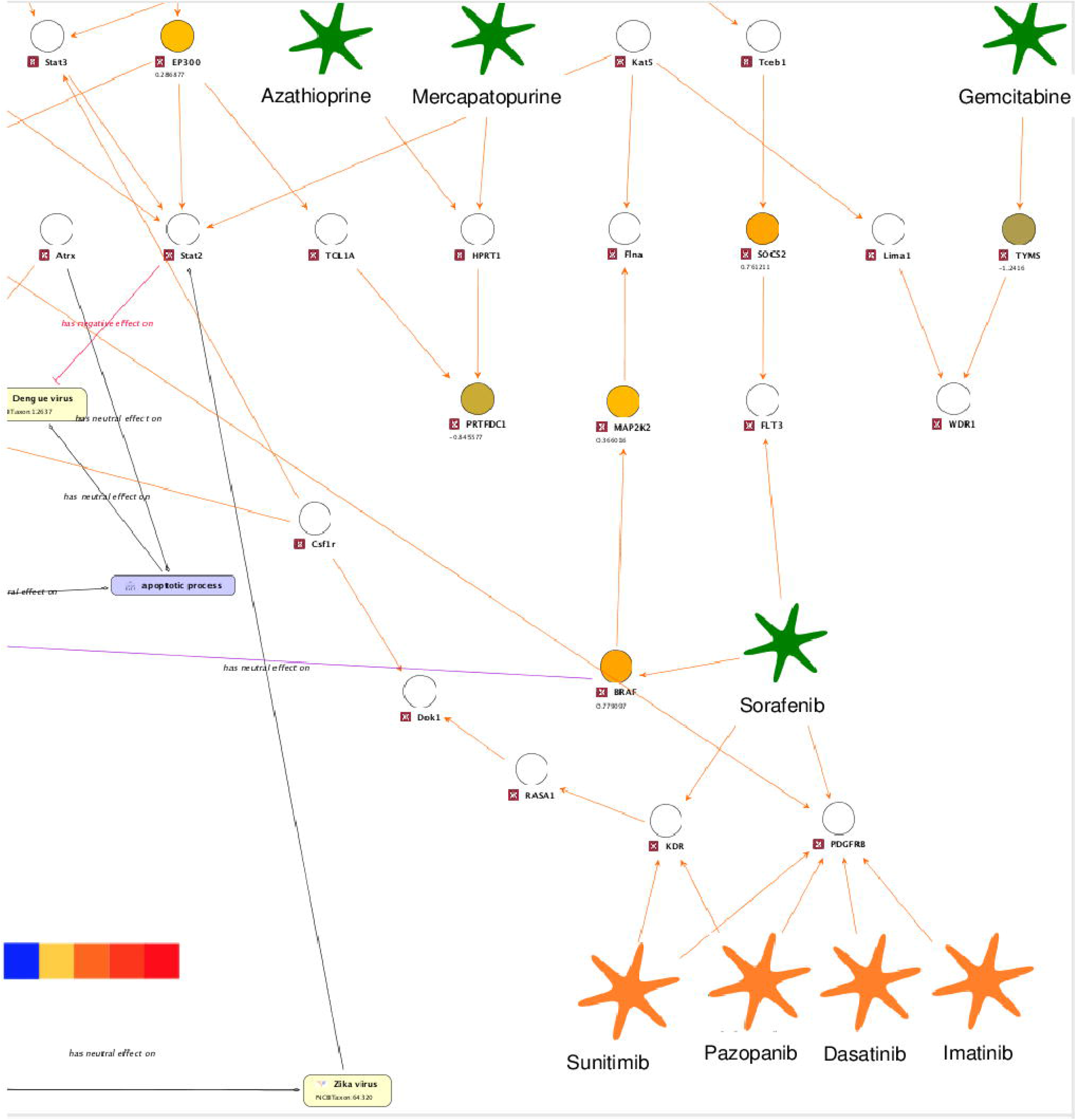
ZIKV molecular interaction and disease map extended for ZIKV effective drugs (enlarged perspective). Up- and down-regulated genes (hNPCs challenged with ZIKV, Tang et al, 2016) are highlighted by a color code ranging from red to blue from enlarged perspectives surrounded by dotted boxes in Figure 7.

The network analysis described above could also be used to rank drugs according to their distance to known ZIKV associated genes, such as STAT2, to suggest a metric for priorisation in screening assays (34). The discussed extended ZIKV map contains 429 target gene products that interact with FDA approved drugs. Evaluation of the distance between STAT2 and drug targets via experimentally proven PPIs revealed that targets of ZIKV effective drugs were on average more proximal to STAT2 compared to other targets.

Combining the proximity measure with additional knowledge, for example the “FDA pregnancy” label, reduces the number of drugs to be screened from 774 to 64.

## Discussion

In case of an emerging epidemic, public health as well as clinical and preclinical research responses are typically hampered by a lack of structured, curated and actionable knowledge. The results of this study describe an approach to knowledge extraction and mapping that can quickly provide an overview of existing and missing information if done by a dedicated and trained group of experts. The developed workflow does not follow formal expert consensus seeking processes, such as Delphi (13), systematic literature review processes such as Cochrane (35) and PRISMA (12), or medical guideline related processes (36,) as these processes are not compatible with the need for speed during emerging epidemics. Nevertheless, the workflow adopts several important aspects of good practice: it is systematic, independent and transparent, provides evidence for all integrated information and uses appropriate quality criteria. Combined with the software tools employed in the process, this pragmatic approach enabled much faster knowledge generation than more traditional methods.

The tools employed in the process need to be able to semantically integrate disparate structured resources of heterogeneous data with ease. However, much of the knowledge that represents scientific research advancements is locked within the unstructured text of classical publications, such as journal articles, newsfeeds or free-form web publications (e.g. Zika-related clinical information at http://www.ovid.com/site/zika/resources.html). The sheer volume of this published information grows constantly and exponentially and, for the most active areas of research, far exceeds the capacity of individual scientists and medical doctors to identify and read all relevant articles. Literature mining, a well-established technology to extract meaningful information from text, provides valuable assistance in structuring the massive amounts of text data and, therefore, is an indispensable tool in the process of guidance generation. Dynamic integration of objects and the relationships they participate in that are present in the literature through the use of structured resources and experimental data is a pre-requisite for analysis and distinguishes the ZIKA KB from text mining-only solutions, such as ContentMine (http://contentmine.org/) or databases dedicated to specific questions such as SncRNAs linking to disease symptoms ((http://zikadb.cpqrr.fiocruz.br/zika/).

Beyond our initial analysis presented here, users can explore the ZIKA KB within a web-based user interface with the use of the online manual (S5) and the step-by-step guide (S4) to reproduce the presented results. We will collect and highly appreciate any user feedback to optimise user experience for broad adoptation. In contrast to alternative useful resources such as the Virus Pathogen Resource (viprbrc.org), which focus on gene and protein sequences, the ZIKA KB integrates genetic, phenotypic and drug knowledge about ZIKV to facilitate the generation of hypotheses, define research priorities and enable better understanding of viral pathogenesis. In addition to interactive exploration, a corresponding ranking of connections in a network based on integration of multiple pieces of biological evidence can also be performed systematically and on a large scale, for example, by applying the ChainRank method (37), which we plan to integrate in the future.

Based on the text mining analyses performed here, disease profiles for the set of five neurotropic flaviviruses were confirmed. Common knowledge was retrieved along with underlying literature evidence and rare manifestations, such as encephalitis, associating with ZIKV and DENV.

Using a molecular interaction and disease map based on ZIKV, Microcephaly, and GBS text mining analyses results, we showed that further exploration of the described map can provide insight into, for example, ZIKV biology, propose conclusions for research decisions, predict drug efficacy, as exemplified in the results section, as well as propose hypotheses on specific host factors and signalling pathways affected by ZIKV. The map can help to distinguish between multiple potential targets of a ZIKV effective drug. The integration of information about effectiveness of other drugs as well as their target genes and the information about genes whose expression is affected during ZIKV infection indicated that Sorafenib likely acts via its target genes, *FLT3* and/or *BRAF*, but not via its alternative target genes *VEGFR* or *PDGFR*. In addition, the number of drugs to be screened was reduced from 774 to 64 by filtering potential drug candidates based on their network distance to ZIKV infection associated genes and additional phenotype relevant additional knowledge, such as contained in the “FDA pregnancy” label.

The conclusions that can be drawn are limited by the initially low number of available publications and limited experimental data, a situation which is inherent to most emerging epidemics. Nevertheless, the work presented shows that the use of a knowledge integrating system can provide guidance for clinical and research responses, such as follow-up studies regarding the association between ZIKV, microcephaly and epilepsy, the validation of candidate drugs for ZIKV treatment, and the validation of candidate genes in specific functional assays to better understand molecular ZIKV infection mechanisms. or to complement existing functional genomic approaches with proteomics studies, such as the integrated proteomics approach identifying cellular targets of ZIKV proteins (38,39). These studies allow additional comparative analyses between ZIKV and other flavivirus family members in terms of virulence and pathogenic traits.

Another limitation of the system is the restricted types of information which can be retrieved by text mining. While qualitative associations between genes/proteins, drugs, diseases and organisms are readily amendable to automatic approaches, it is currently almost impossible to extract, for example, clinical study designs, detailed quantitative information or complex treatment plans.

Finally, the ZIKA KB in its current stage enables exploration of the integrated information, as well as generation and curation of text-mining analysis but is not a public tool for molecular interaction and disease map generation. The functions required for these tasks will need further refinement before they can be made available in a general way.

In summary, this approach in our opinion, provides a feasible way to collect and integrate existing knowledge to better understand the molecular mechanisms of an emerging pathogen. In addition our approach helps to identify gaps in knowledge and, together with the other features, guides rapid and effective responses to future epidemics. We have made the specific outcome of our approach, the Zika KnowledgeBase, publicly available as a hopefully valuable resource to the ZIKV research community.

In the light of the current COVID-19 pandemic we now apply the described workflow to SARS-CoV-2 and other coronaviruses and will make the developed resource available as described.

## Acknowledgements

The authors wish to thank their colleagues for discussion and feedback, and Sheridon Sauer and Shannon Frances for critical review of the article.

## Supporting Information

**S1 Fig. Molecular interaction and disease map generation protocol**

**S2 Fig. ZIKV KnowledgeBase Data Model**

Based on input from clinical and virology experts the concepts required to represent existing pathophysiological knowledge of infectious diseases were modelled using the BioXM Knowledge management environment. To this end objects (as nodes) and relations (as edges) are defined on an abstract level such as “gene”, “disease” or “interacts with”. For the ZIKV KB we focused on concepts required to represent text mining results and information from structured databases of protein-protein and drug – protein interaction, namely genes, diseases, pathogens and drugs.

Pathogens are displayed as yellow nodes and were uniquely identified by using the NCBI Taxonomy ontology. Compounds (white nodes) were referenced to entries of the ChEBI ontology. Diseases (violet nodes) and genes (orange nodes) can reference one or multiple sources such as ICD10, Entrez or Ensembl. Relationships between pathogens, genes, drugs and compounds were defined by three types of text mining relations, upregulation (green edges), downregulation (red edges) and regulation (black edges). Associated information, not represented as object or relation, is being stored as so called “annotation” basically a note that can be assigned to any semantic concept. The annotation form “Textmining information evidence”, for example, stores information, such as the sentence containing the extracted statement (field “Reference”) or the predicate (field “Interaction type”) and links directly to the PubMed entry (represented as BioRS entry: MEDLINE). To enable experts to support or contradict extracted text mining relationships the annotation form “Textmining Validation” was introduced to generate a configurable curation workflow. Molecular interaction and disease maps are modelled using the “context” object type to group specific sub-networks (interactions between genes, diseases, pathogens and drugs) within the global semantic network. PPI and drug-protein interactions are represented by the “Interaction” relation (orange edges) with supportive evidence stored in two annotation forms (“BioGRID Interaction Evidence” and “Interaction Information” storing information from the VirHostNet database), and can be added as additional sub-networks to a disease map.

Based on the data model dynamic visualisation can be generated, for example drug targets or differentially expressed genes based on configurable queries and views. The data model is automatically transformed into a natural language like query and reporting language. This language can be used to e.g. define a query retrieving the number of drugs that interact with a protein of interest, and generating a view which applies a colour code to a gene according to the number of interacting drugs.

**S3 Fig. Molecular interaction and disease map wizard.**(i) A text mining analysis result (or multiple ones) can be chosen from to start populating the map. (ii) Based on the genes extracted from text mining relationships (highlighted in orange, top left) a network search algorithm is applied to extend the map with PPI & protein-drug interactions. (iii) Upon selection of wanted interaction data the final map is automatically rendered in different perspectives, displaying literature evidence, experimental data or drug targets (shown from left to right, bottom).

**S4 Step-by-step guide to reproduce the results in the web interface**

**S5 Manual for web interface**

