## Supporting Information S4 for "Informing epidemic (research) responses in a timely fashion by knowledge management - a Zika virus use case"

To reproduce the uses and use cases described in the manuscript please register yourself at https://ssl.biomax.de/zikakb/ and login to the Zika Knowledge Portal. There follow the step-by-step description below which references the corresponding sections of the Zika KB manual (Supplement 2 or online at the Zika Knowledge Portal).

**1. Get an overview of ZIKV biology**

Diseases associated with Zika infections can be accessed by steps 2.3.2 "Flavivirus Diseases" and 2.3.3 "Disease-disease associations" of the manual.

Zika virus specific interactions with host genes are available by step 2.3.4 "Zika Virus Host Genes" of the manual.

Integrated literature mining results, database content and expert knowledge are available with step 2.4 "disease maps" of the manual.

**2. Predict drug efficacy**

Step 2.3.6 "ZIKV effective drugs" of the manual shows how to check for drugs interacting with host genes associated with Zika infection.

**3. Indicate specific host factors/signalling pathways affected by ZIKV**

The steps described in 2.3.4 "Zika host gene interaction" and 2.3.5 "Flavivirus host gene interaction" allow to query for genes and pathways associated with Zika or Flavivirus infection.

**4. Generate hypothesis**

We used steps 2.3.2 "Flavivirus Diseases", 2.3.3 "Disease-disease associations", 2.3.4 "Zika Virus Host Genes" and 2.3.5 "Flavivirus host gene interaction" to generate hypothesis about differences between Zika biology and that of other Flaviviruses.

**5. Share knowledge**

Currently knowledge is shared from PREPARE experts to the Zika community via 2.4 "Disease maps". Contribution to the curation process is open by contacting the authors.

**6. Identify knowledge gaps**

By executing steps 2.3.7 “Studies with flavivirus challenge” and 2.3.8 “Studies with virus challenge (neuro cell)” of the manual it is possible to quickly get an overview of existing studies and the gaps left open by them.

**7. Explore data**

Integrated experimental data can be explored at steps 2.3.2 “Flavivirus diseases”, 2.3.3 “Disease-disease association”, 2.5 VirHostNet Interactions and 2.6 “Virus-Host interactions”.

**8. Interpret data**

Interpretation of integrated experimental data in the light of knowledge about Zika and host biology was performed by executing the tasks described in 2.3.4 “Zika Virus Host Genes”, 2.3.5 “Flavivirus Host Genes” and 2.3.6 “ZIKV effective drugs”.

**9. Computational data analysis**

Computational analysis of experimental data is greatly improved by incorporating existing, integrated knowledge. These are available from the Zika KB by exporting relevant integrated information for example at steps 2 “Zika Virus Host Genes” and 2.3.5 “Flavivirus Host Genes” regarding host genes and their functions or 2.3.4 “Disease maps” and 2.5/2.6 “Virus-Host Interactions” regarding interactions between Zika and host.

**10. Predictive model generation**

To generate predictive computational models either a statistical or a knowledge based approach can be followed.
The statistical approach benefits from incorporating existing, integrated knowledge in the same as the computational data analysis described above by exporting relevant integrated information for example at steps 2 “Zika Virus Host Genes”, 2.3.5 “Flavivirus Host Genes” regarding host genes and their functions or 2.3.4 “Disease maps” and 2.5/2.6 “Virus-Host Interactions” regarding interactions between Zika and host.

The knowledge based approach directly depends on expert validated, structured descriptions of biological processes as provided by step 2.4 “Disease maps”.

**11. Visualise drug targets and host factors involved in ZIKV pathogenesis**

Step 2.3.6 “ZIKV effective drugs” of the manual directly describes how to visualise drug targets and host factors involved in Zika virus pathogenesis.

**12. Interrogate network to identify host factors targeted by virus/interact with drugs**

Step 2.3.6 “ZIKV effective drugs” of the manual directly describes how to identify host factors that interact with a virus and/or are targeted by drugs.

**13. Explore network by overlaying expression data (RNASeq)**

Step 2.3.6 “ZIKV effective drugs” of the manual directly describes how to visualise RNASeq data on top of a virus – host interaction network.

**14. Explore integrated literature for relevance (by graph information layer) -> Fig 4a**

Step 2.4.1 “Disease map report” of the manual directly describes how to access the network display of a disease map. Clicking on “ZIKV” within in the ZikaBKB Disease Maps list redirects to the Disease map report. By using the “Graph” action menu on top of the page the network of ZIKV disease map is displayed. On the upper left of the network select “Literature evidence” (default is “Disease map ZikaKB”) to display the amount of literature evidence as highlighted by the thickness of the edges.

**15. Overview on drug targets**

Step 2.3.6 “ZIKV effective drugs” of the manual directly describes how to visualise drug targeting information on top of a virus – host interaction network.

**16. Explore critical genes within gene - disease – pathology networks**

Step 2.3.6 “ZIKV effective drugs” of the manual directly describes how to visualise complex information on top of the pathology network and thereby identify genes of importance.

**17. Disease associated with virus of interest**

To identify disease associated with a Flavivirus of your interest please follow steps 2.3.2 “Flavivirus diseases”, 2.3.3 “Disease-disease association” of the manual.
