## Supporting Information S5 for "Informing epidemic (research) responses in a timely fashion by knowledge management - a Zika virus use case"

### Zika Knowledge Base
### User Guide

**2020**

Biomax Informatics AG
Robert-Koch-Str. 2
82152 Planegg
Germany

Website: www.biomax.com

*Zika Knowledge Base User Guide*

*BioXM version 6.2 build 263, December 2019*

*© 2019 Biomax Informatics AG, all rights reserved*

*This document and the product it describes are subject to change without prior notice. This document does not represent a commitment on the part of Biomax Informatics AG or its distributors. Biomax makes no representations or warranties with respect to the contents hereof and specifically disclaims any implied warranties of merchantability or fitness for any particular purpose.*

*The software described in this document is furnished under a license agreement. The software may be used or copied only in accordance with the terms and conditions of the license agreement. It is against the law to copy the software on any medium except as specifically allowed in the license agreement.*

*No part of this document may be reproduced or transmitted in any form or by any means, whether electronic or mechanical, including photocopying, recording or information recording and retrieval systems, for any purpose other than the purchaser's personal use, without the prior express written permission of Biomax.*

*Biomax Informatics AG; Sitz: Martinsried bei München, Amtsgericht München, HRB 134442; Ust-IdNr. DE 191 603 142; Steuer-Nr.: 143 / 100 / 10945; Vorstand: Dr. Klaus Heumann (Vorsitzender), Dr. Hein Paul Osenberg; Vorsitzender des Aufsichtsrates: Prof. Dr. Hans-Werner Mewes*

*Biomax, BioRS, and BioXM are registered trademarks of Biomax Informatics AG in Germany and other countries. Registered names, trademarks, etc., used in this document, even when not specifically marked as such, are not to be considered unprotected by law.*

Help for working with BioXM knowledge portals in general

This section covers working with BioXM knowledge portals in general. Because the interface is highly flexible, your knowledge base may differ. Specific information about your knowledge base is available in Section 2 Help for the Zika knowledge base

#### 1.1. Logging in

Complete the following steps to login into a BioXM web portal:

1. Open a browser (Google Chrome®, Firefox® or Safari® browsers are recommended) and enter the uniform resource locator (URL) for the corresponding BioXM web portal (https://ssl.biomax.de/zikakb). The login page will be displayed.
2. Enter your user name and password.
3. Click the "Login" button.

The BioXM web portal homepage will be displayed.

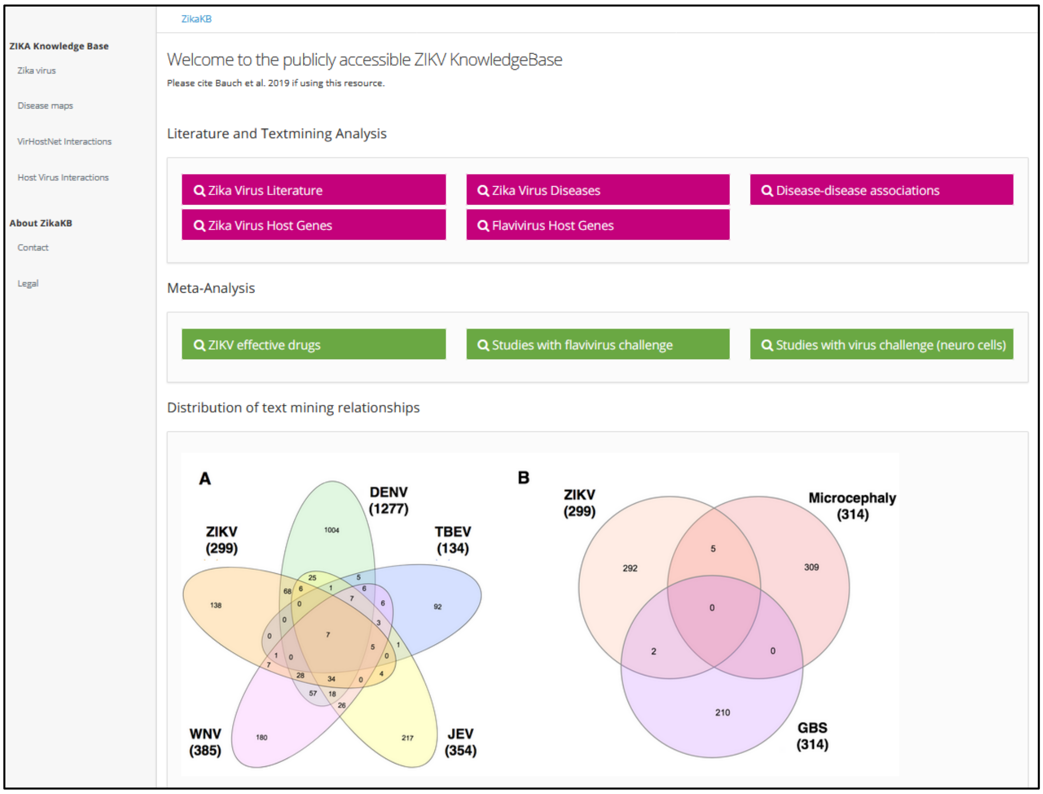

**Homepage of the Zika knowledge base portal**

#### 1.2. Navigating the interface

The BioXM web portal interface provides access to all content, resources and tools available in your installation. The interface is highly flexible, but in general, can be divided into three areas:

- **Navigation menu** (on the left) provides links to content, resources and result sets available in your installation.

- **Content panel** (the main part of the window) contains the content, resource or result set that has been selected. In addition, there may be additional navigation items. The blue text items, buttons and icons are, in most cases, links to related information.
- **Toolbar** (near the upper right corner) provides additional buttons to general actions, such as printing the current page or logging out. Click one of the buttons to perform the desired action

#### 1.3. Using search forms

In many areas, search forms are available for starting a query or search. Specific instructions are available in the individual search forms. In general, complete the following steps to start a search:

1. Enter search term(s) in the field(s). To see all results, enter an asterisk (*) in the field.
2. Click the "Search" button.

The search results (result set) will be displayed. The search can be refined directly in the result set (see Section 1.5 Searching in tables/result sets).

Click the "Clear" button to clear the search fields.

#### 1.4. Working with tables

Search results and other information are often displayed in a table. This section describes working with tables in the BioXM system in general. The actions available in individual tables may vary.

In most tables, several actions can be performed directly:

- **Display all columns** (potentially hidden on small screens e.g. mobile devices) by clicking the “+” sign at the beginning of the row
- **Display more information about the entry** by clicking a link in the table.
- **Sort the results** by clicking the table headers in any column.
- **Change the columns shown in the table** by clicking the right arrow icon in the upper left corner of the table. In the dialog, check the box of the columns to be displayed.
- **Search the table contents** using the fields above the table and the "Search" button. See Section 1.5 Searching in tables/result sets for more information.
- **Change the number of entries displayed per page** using the "Rows" drop-down menu above the table.
- **Scroll through additional pages** using the arrow buttons below the table or entering a number in the "Page" field.
- **Export table contents** to an Excel table or in tab-delimited format using the "Export" drop-down menu above the table. See Section 1.6.1 Exporting results from a table for more information.

#### 1.5. Searching in tables/result sets

Table contents can be searched. In this way, a search on a result set can be performed or refined.

1. Select the column in which to search from the drop-down menu above the table.
2. Enter the search term in the field next to the "Search" button.
3. Click the "+" (plus) button to add additional search terms and columns.
4. Click the "Search" button.

5. The search results will be displayed in a table.

#### 1.6. Exporting

##### 1.6.1. Exporting results from a table

Table contents can be exported to an Excel table or in tab-delimited format using the "Export" drop-down menu above the table.

1. Select the rows to export by putting a check mark in the first column. (You can select and de-select all rows by clicking the check mark in the header row.)
2. Select the export method ("Excel" or "Tab-delimited") in the "Export" drop-down menu above the table.
3. A progress dialog will be displayed. For larger exports there is the option to put the export to the background. When the export is finished, you can get the exported file by clicking the "Get file" button.

##### 1.6.2. Retrieving an export put in the background

To view an export put into the background, click the "Background exports" option in the "Export" drop-down menu above the table. A list of the exports which have been put into the background will be shown. Select the export you wish to retrieve.

#### 1.7. Working with panels

### 1.7.1.

Reports may use multiple panels to structure a page. Every panel offers three interactions

1. “-/+“ icon – collapse/expand the panel to hide/access its content
2. Double arrow icon – expand/collapse panel to/from full screen

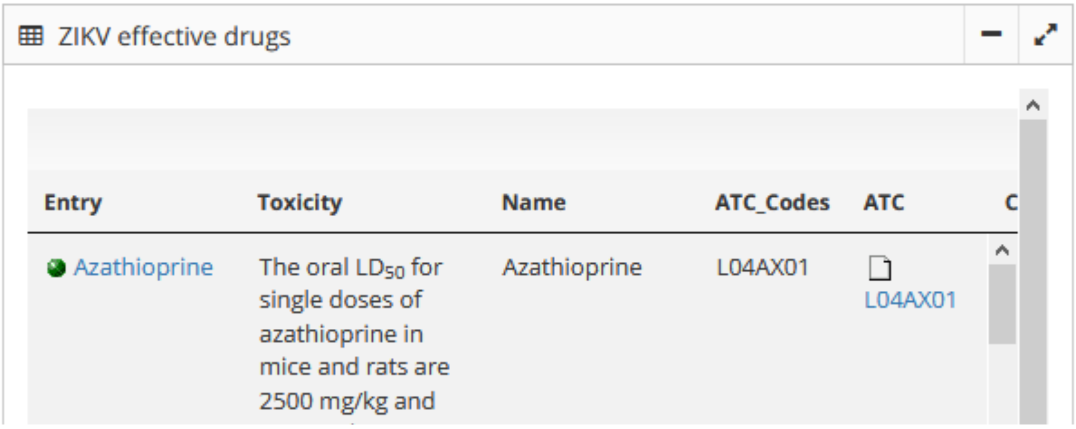

#### 1.8. Working with graphs

Access to integrated knowledge networks is often provided in graph form. Every graph provides the following actions:

3. Paper sheet icon – display the graph objects as table report

4. Rhombus icon – zoom to fit the graph into the available window
5. Magnifier "+" icon – zoom in
6. Magnifier "-" icon – zoom out
7. Drop down list – select from predefined information overlays
8. Arrow icon – Export to external graph viewer

Clicking on an individual item (node or edge) in the graph will open a detailed report of it.

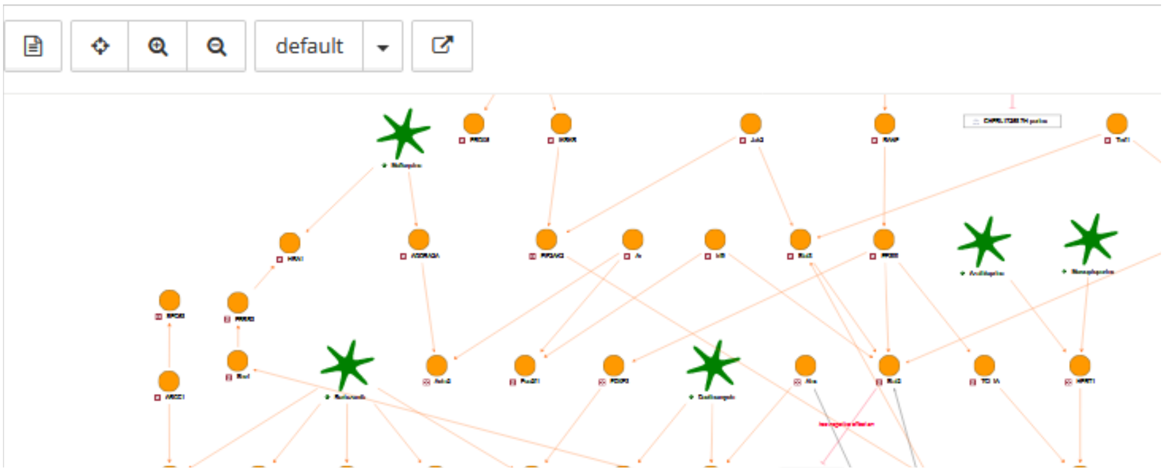

### 2. Help for the Zika knowledge base

#### 2.1. Home

Upon logging in, the knowledge base homepage is displayed. It can be displayed at any time by clicking the "Zika KB" link in the top bar.

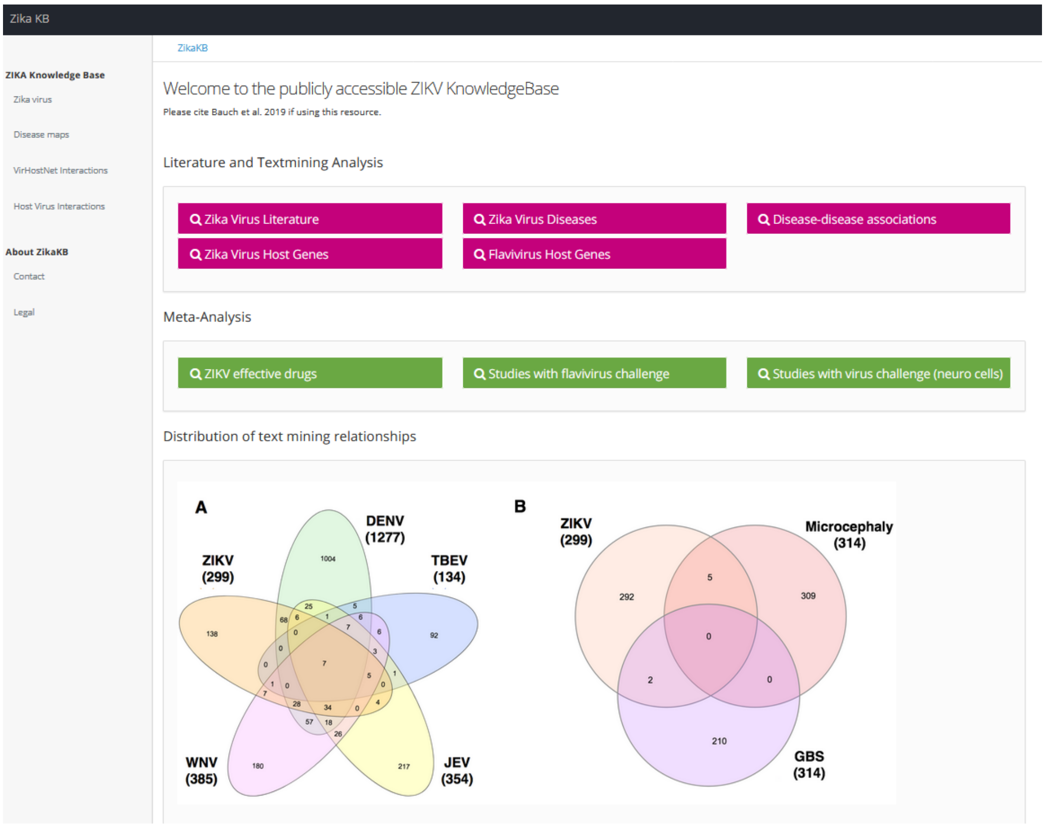

**Homepage of the Zika knowledge base portal**

The homepage provides access to all content in the knowledge base. Click an icon or link to navigate to the desired section.

##### 2.1.1. General search

The following options are available:

- Search and Browse Zika related information — described in Section 2.3
- Browse manually curated Disease maps —described in Section 2.4
- Search Virus Host molecular interactions from VirHostNet – described in Section 2.6
- Search Host Virus Interactions curated by PREPARE from literature - described in Section 2.7
- Literature mining — described in Section 2.7

#### 2.2. Help

Online help for the knowledge base can be displayed by clicking the **Manual** link in the left menu.

#### 2.3. Zika virus

The **Zika virus** page provides a number of Zika related searches which are accessible by individual buttons and are described in detail below.

##### 2.3.1. Zika Virus Literature

Select the **Zika KB** link in the top bar and then the pink **Zika Virus Literature** button.

Searches Medline for any Zika related publication using the BioRS plugin and displays the list of Medline abstracts currently associated with Zika.

The results are displayed as a table. The table can be used as described in Section 1.4 Working with tables.

The columns are described below:

1. PMID — PubMed ID linked to the full Medline entry
2. Full text — if manually uploaded by a curator the full text of article is displayed
3. Authors — list of author names
4. Title – publication title as provided by Medline
5. Journal — name of the publication source
6. Year — publication year
7. DB – for sequence related publications Medline provides information about the corresponding sequence database
8. Curation notes – if manual curation occurred the curators notes are displayed
9. Last curator – name of the last curator working on the entry

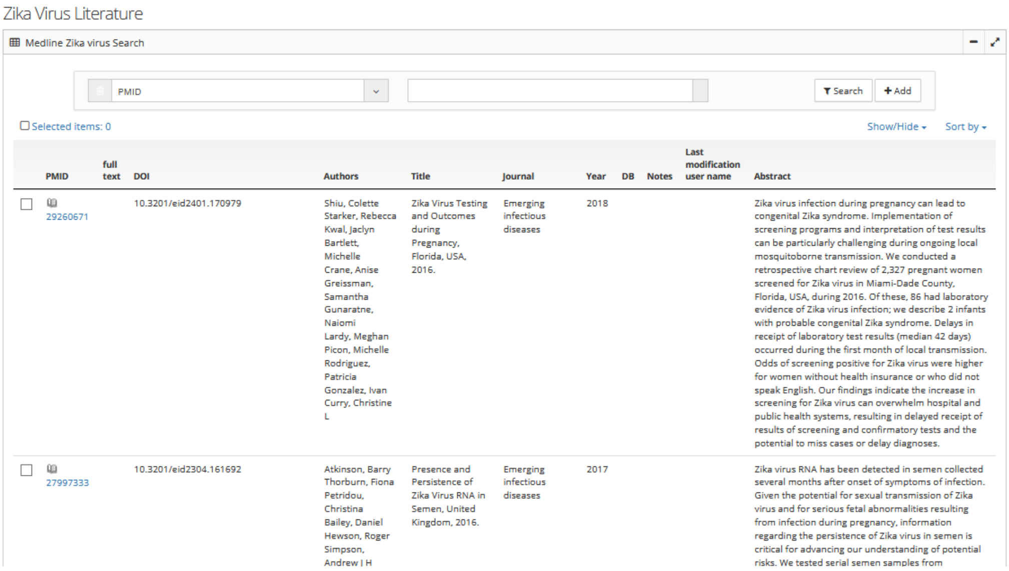

Zika Virus Literature

Medline Zika virus Search

PMID | Search | Add

Selected items: 0 | Show/Hide | Sort by

| PMID | full text | DOI | Authors | Title | Journal | Year | DB | Notes | Last modification user name | Abstract |
|---|---|---|---|---|---|---|---|---|---|---|
| 29260671 | | 10.3201/eid2401.170979 | Shiu, Colette Starker, Rebecca Kwal, Jaclyn Bartlett, Michelle Crane, Anise Greissman, Samantha Gunaratne, Naiomi Lardy, Meghan Picon, Michelle Rodriguez, Patricia Gonzalez, Ivan Curry, Christine L | Zika Virus Testing and Outcomes during Pregnancy, Florida, USA, 2016. | Emerging infectious diseases | 2018 | | | | Zika virus infection during pregnancy can lead to congenital Zika syndrome. Implementation of screening programs and interpretation of test results can be particularly challenging during ongoing local mosquitoborne transmission. We conducted a retrospective chart review of 2,327 pregnant women screened for Zika virus in Miami-Dade County, Florida, USA, during 2016. Of these, 86 had laboratory evidence of Zika virus infection; we describe 2 infants with probable congenital Zika syndrome. Delays in receipt of laboratory test results (median 42 days) occurred during the first month of local transmission. Odds of screening positive for Zika virus were higher for women without health insurance or who did not speak English. Our findings indicate the increase in screening for Zika virus can overwhelm hospital and public health systems, resulting in delayed receipt of results of screening and confirmatory tests and the potential to miss cases or delay diagnoses. |
| 27997333 | | 10.3201/eid2304.161692 | Atkinson, Barry Thorburn, Fiona Petridou, Christina Bailey, Daniel Hewson, Roger Simpson, Andrew J H | Presence and Persistence of Zika Virus RNA in Semen, United Kingdom, 2016. | Emerging infectious diseases | 2017 | | | | Zika virus RNA has been detected in semen collected several months after onset of symptoms of infection. Given the potential for sexual transmission of Zika virus and for serious fetal abnormalities resulting from infection during pregnancy, information regarding the persistence of Zika virus in semen is critical for advancing our understanding of potential risks. We tested serial semen samples from |

**Zika Virus Literature -** Medline search results table

##### 2.3.2. Flavivirus Diseases

Select the **Zika KB** link in the top bar and then the pink **Flavivirus Diseases** button.

Searches stored literature mining results for occurrence of any disease with a given selection of virus e.g. Zika, Flavivirus.

To illustrative the purpose of this function the page is headed by an overview of the association of different Flavivirus with different diseases reflecting the status published in Bauch et al. Figure 5

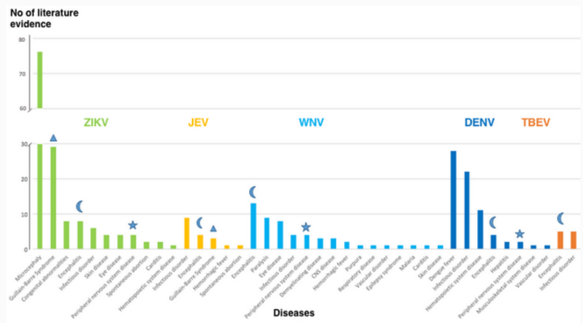

The number of literature evidence (y-axis) for each of the **diseases** (x-axis) is grouped for the five flaviviruses. Symbols highlight neurological diseases associated with more than 1 virus.
Triangle: GBS, moon: Encephalitis, pie-chart: Peripheral nervous system disease, star: Neurologic manifestations
**Diseases** displayed represent a slim version of the Human Disease Ontology

To perform your own search:

1. Select the flavivirus(es) of interest from the top search field drop-down list (Virus entries)
2. Select (or de-select, by clicking on the “x”) the literature mining of interest from the “Literature mining analysis” search field drop-down list
3. Click the "Search" button to perform the search.

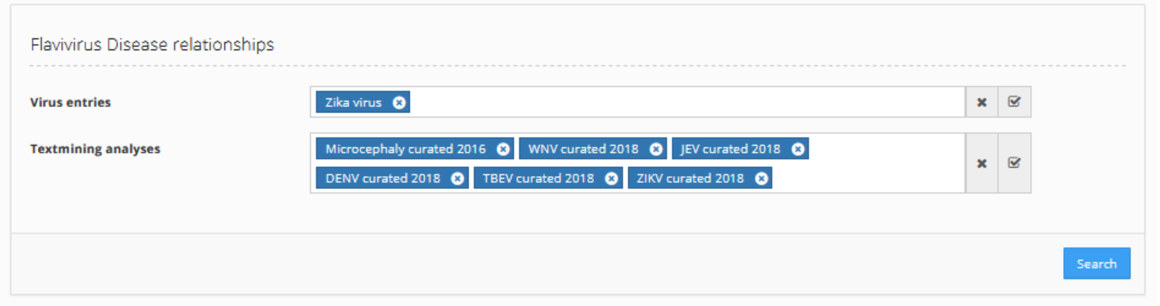

The results will be displayed as a table. The table can be used as described in Section 1.4 Working with tables.

The columns are described below:

1. Disease — the disease associated by literature mining with the virus as mapped to the standard vocabularies used within the ZikaKB
2. UID — unique identifier it the disease retrieved by literature mining was directly mapped to an existing disease ontology
3. DisGenNet — genes associated by the DisGenNet database to the given disease
4. Synonymous NCI Thesaurus – diseases synonyms provided by the NCI Thesaurus as mapped to the ZikaKB standard vocabulary
5. Synonymous DO — diseases synonyms provided by the Disease Ontology as mapped to the ZikaKB standard vocabulary
6. DO — higher level Disease Ontology entry mapped to the literature mining mapped disease entry
7. Parent Disease – higher level NCI Thesaurus entry mapped to the literature mining mapped disease entry
8. Synonyms – any synonyms mapped to the standard vocabulary used within the ZikaKB
9. No of evidences – number of specific virus – disease associations retrieved by literature mining
10. Predicate – type of virus – disease association retrieved by literature mining e.g. “associates”, “causes”, “mutation”
11. Lit Source – PubMed ID or full text entry from which the virus – disease association was retrieved
12. Source – association source, in this case either the specific virus or the disease
13. Target – association target, in this case either the specific virus or the disease
14. Relation target type – type of the association, in this case either “Disease” or “Taxonomic”
15. Literature mining – history of individual literature mining runs which retrieve the corresponding association

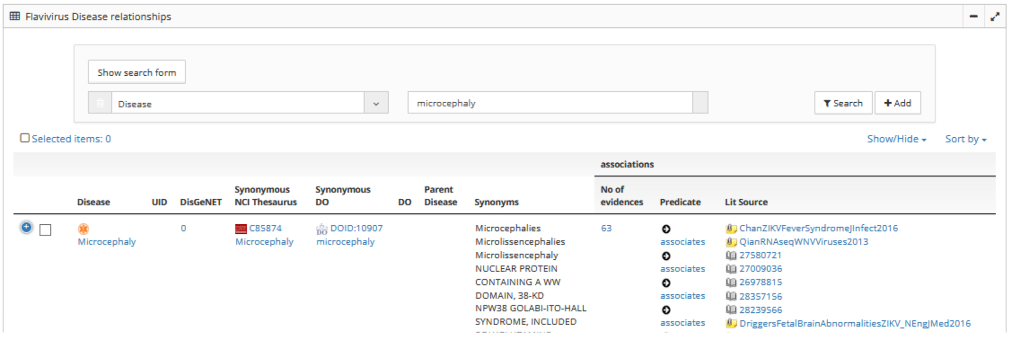

**Zika Virus Disease** –The table of interactively retrieved Zika – disease associations reflecting the selected snapshot of literature mining (here 2018). Note the “+” Button in front of the first row to display further columns as described above.

##### 2.3.3. Disease-disease associations

Select the **Zika KB** link in the top bar and then the pink **Disease-disease associations** button.

Searches stored literature mining results for co-occurrence of any disease and the virus this co-occurrence is associated with.

To illustrative the purpose of this function the page is headed by an overview of the association of different disease co-occurrences using the Dengue virus literature mining and reflecting the status published in Bauch et al.

| **Comorbidity** | **PMID** | **Source** | **Target** | **Predicate** | **Infection** |
|---|---|---|---|---|---|
| Encephalitis | 23487229;9266146 | Dengue Fever | Encephalitis | associates | DENV |
| Glucose intolerance | 20639831 | Glucose intolerance | Dengue disease | associates | DENV |
| Guillain-Barre syndrome | 26895226;21695889 | Dengue disease | Guillain-Barre syndrome | associates | DENV |
| Hematopoietic system disease | 26755235;22828377;19237145;1511177 | Dengue disease;Hepatitis | Disseminated intravascular coagulation;Aplastic anemia;Aplasia | asociates;induces | DENV |
| Hydrocephalus | 24059883 | Dengue disease | Hydrocephalus | associates | DENV |
| Lupus erythematosus | 24204176 | Dengue disease | Lupus erythematosus | induces | DENV |
| Respiratory disease | 8687212; 24825269;8687212 | Dengue disease | Adult Respiratory Distress Syndrome Adverse Event | associates | DENV |
| Uveitis | 26229512 | Dengue disease | Panuveitis | associates | DENV |
| Vascular Disorder | 24772398;25884693;23785539 | Dengue disease | Hypotension;Cardiovascular System Finding;Hypovolemic Shock | associates;involves;leads to | DENV |
| Movement disease | 16292498 | Movement disease | Japanese encephalitis | associates | JEV |
| Vascular Disorder | 15650547 | Japanese encephalitis | Cerebral Infarction | associates | JEV |
| Vascular Disorder | 23514435 | Branch retinal artery occlusion | Posterior Uveitis | associates | WNV |
| Musculoskeletal system disease | 27229696 | Rhabdomyolysis | Myositis | associates | ZIKV |

To perform your own search:

1. Select the literature mining of interest from the “Literature mining analysis” search field drop-down list
2. Click the "Search" button to perform the search.

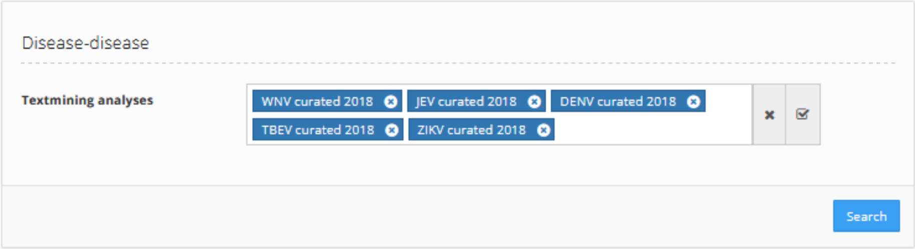

The results will be displayed as a table. The table can be used as described in Section 1.4 Working with tables.

The columns are identical to those described in 2.3.2 with the only difference that all disease information’s are occurring twice, for the first disease and its co-morbidity.

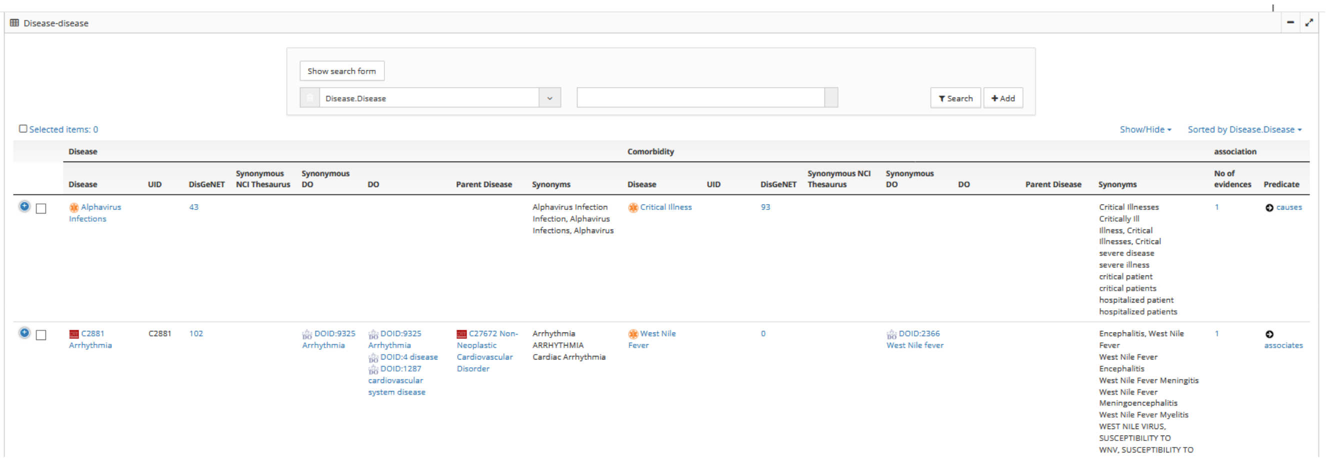

**Disease – disease association** –The table of interactively retrieved Disease–disease associations reflecting the selected snapshot of literature mining (here 2018). Note the “+” Button in front of the first row to display further columns as described above.

##### 2.3.4. Zika Virus Host Genes

Select the **Zika KB** link in the top bar and then the pink **Zika Virus Host Genes** button.

Searches existing literature mining results for host genes interacting with Zika virus genes and displays the list of host genes currently associated with Zika.

The result page provides two panels:

The left panel shows a summary for all interacting host genes in terms of their involvement in signal transduction pathways (using KEGG gene annotations) and biological processes (using GO gene annotations). Selecting any of the bars in the histogram redirects to the list of genes visualized within the selected bar. From this page the list of genes can be selected for Export to Excel using the Export button in the top menu bar. Clicking the Browser “Back” button redirects to the **Zika Virus Host Genes** page.

The right panel provides a list of host genes found to interact with Zika in tabular form.

The columns are described below:

1. Gene – HGNC gene name
2. GO slim — localization of host gene as derived from a reduced GO ontology
3. Biological Process – functional involvement of host gene derived from GO annotations
4. KEGG – signal transduction pathway the host gene is associated with derived from KEGG annoatations
5. Synonyms – synonyms of the host gene used for literature mining
6. No of evidences – number of publications that provide information about the specific host gene – Zika association
7. Predicate – type of host gene – Zika interaction

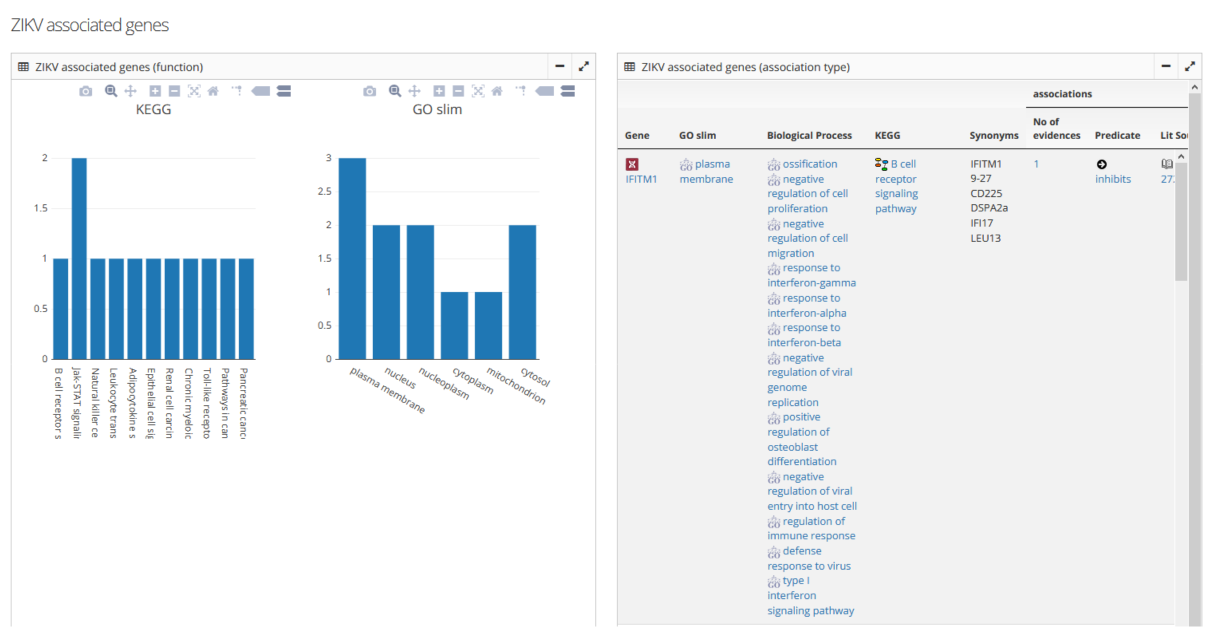

**Zika Virus Host Gene** –Left panel providing a graphical overview on host gene involvement in signal transduction pathways and biological processes. Right panel lists the individual interacting host genes with interaction type, functions and number of evidences.

##### 2.3.5. Flavivirus Host Genes

Select the **Zika KB** link in the top bar and then the pink **Flavivirus Host Genes** button.

Searches existing literature mining results for host genes interacting with a predefined list of Flaviviruses and displays the list of host genes currently associated with ZIKV.

The result page provides an individual panel for each of the selected Flaviviruses (ZIKV, DENV, WNV, and JEV). Each panel contains a graphical summary of the biological function and a list of host genes as explained in 2.3.4.

Selecting any of the bars in the histogram redirects to the list of genes visualized within the selected bar. From this page the list of genes can be selected for Export to Excel using the Export button in the top menu bar. Clicking the Browser “Back” button redirects to the **Flavivirus Host Genes** page.

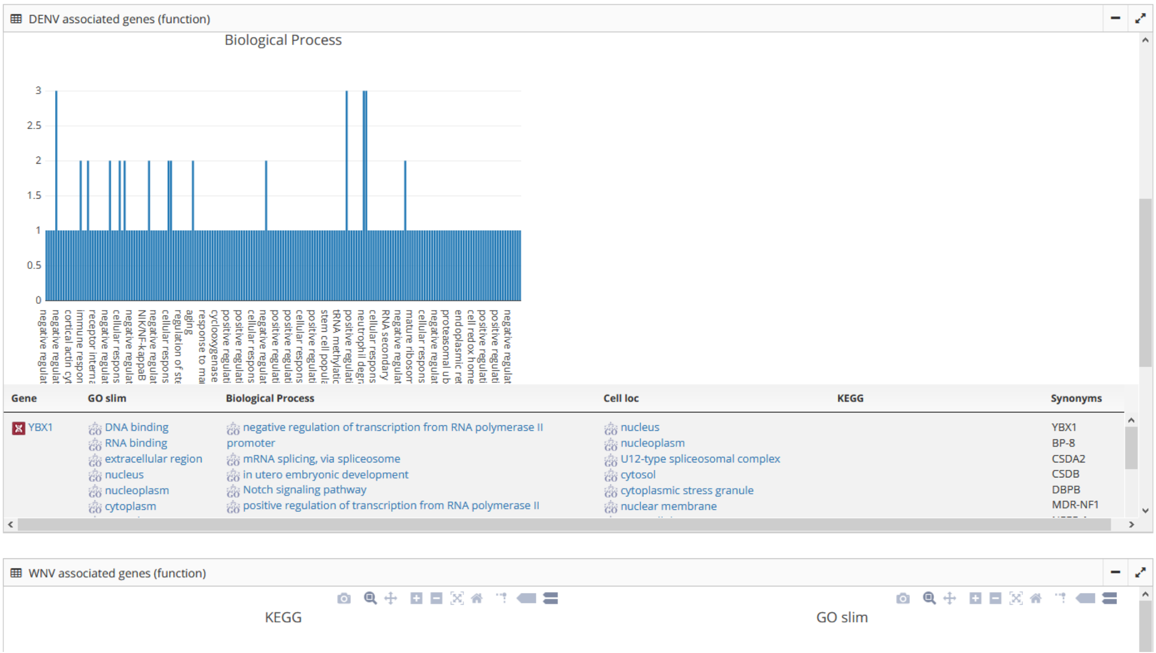

**Flavivirus Host Gene** –Upper part of each panel providing a graphical overview on host gene involvement in signal transduction pathways and biological processes. Lower part listing the individual interacting host genes with interaction type, functions and synonyms.

##### 2.3.6. ZIKV effective drugs

Select the **Zika KB** link in the top bar and then the green **ZIKV effective drugs** button.

Based on the Zika virus host gene interactions described in 2.3.4 the drugs interacting with those host genes are searched.

A list of drugs interacting with currently known host genes associated with Zika is displayed.

The result page provides two panels.

The upper panel displays a graphical network view of Zika and interacting host genes as well as FDA approved drugs found to associate with these host genes. The graph dropdown list allows to overlay the network with

1. Drug targets – Colour coding of genes based on the number of any drug associated with them in DrugBank
2. hNPC ZIKV — Colour coding of host genes according to their expression change in publicly available Zika infection assay transcriptomics measurements (Tang et al. 2016)

The lower panel displays the list of FDA approved drugs that associate with host genes interacting with Zika.

The columns are described below:

1. Entry – drug name
2. Toxicity – available toxicity information
3. Name – other drug names
4. ATC – drug Antomical Therapeutic Chemical Classification
5. Pregnancy category

a. Category A: Adequate and well-controlled studies have failed to demonstrate a risk to the fetus in the first trimester of pregnancy

b. Category B: Animal reproduction studies have failed to demonstrate a risk to the fetus and there are no adequate and well-controlled studies in pregnant women

c. Category C: Animal reproduction studies have shown an adverse effect on the fetus, and there are no adequate and well-controlled studies in humans, but potential benefits may warrant use of drug in pregnant women despite potential risks

d. Category D: There is positive evidence of human fetal risk based on adverse reaction data from investigational or marketing experience or studies in humans, but potential benefits may warrant use of drug in pregnant women despite potential risks

6. ZIKV effective drug - cell line used for assessment of drug (HuH-7, HeLa, JEG3, hNSC, HAEC)
7. Drug target – Gene targeted by drug and if available change in expression
8. FDA – FDA status of drug
9. Type – drug type e.g. small molecule, biosimilar
10. Description – description of drug e.g. use, mode of action
11. Drug category – category of drug e.g. steroid
12. Chemical formula – chemical formula of drug
13. IUPAC name – drug IUPAC name
14. Drug targets – target type of the drug e.g. topoisomerase
15. Synonyms – synonymous names used for the drug
16. PDB ID – PDB ID of proteins targeted by drug
17. PubChem – PubChem ID of drug
18. UniProt – UniProt ID of proteins targeted by drug
19. Drug reference – publications associated with the drug
20. PMID – PubMed ID of publications associated with the drug

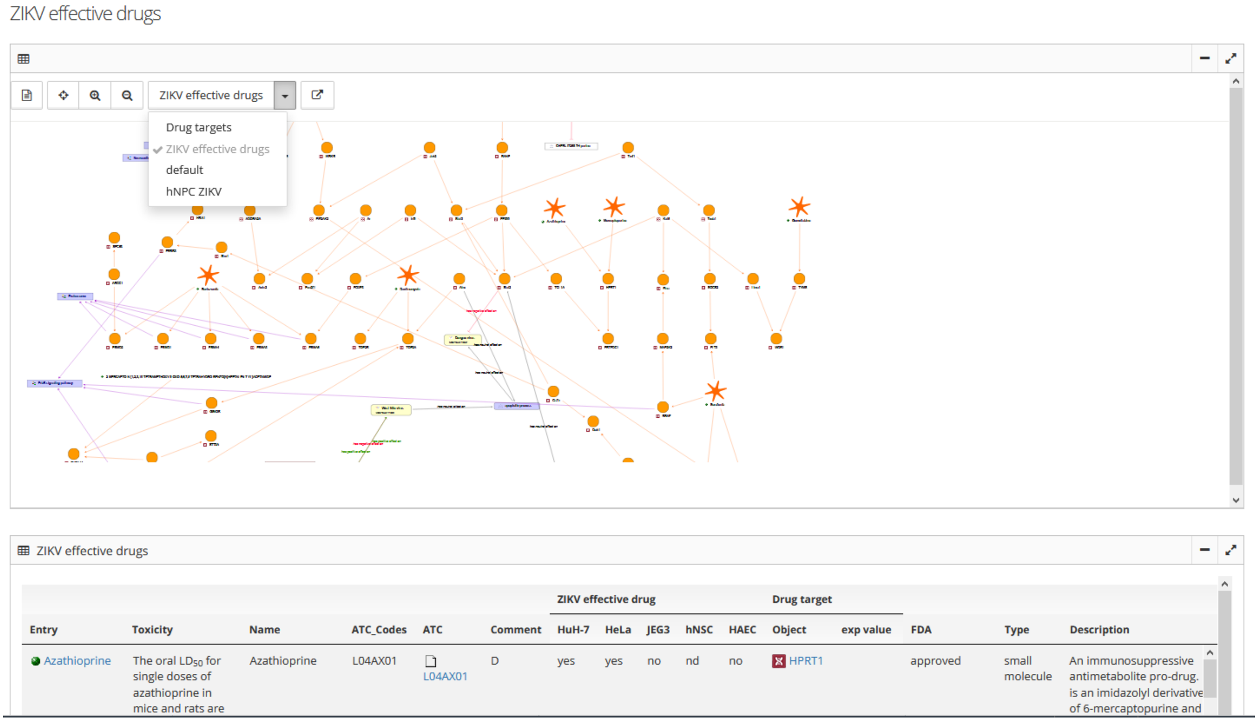

**ZIKV effective drugs** –Upper panel providing a graphical network of drugs associated with host genes interacting with Zika. Lower part listing the individual drug with detailed information.

##### 2.3.7. Studies with flavivirus challenge

Select the **Zika KB** link in the top bar and then the green **Study with flavivirus challenge** button.

Studies providing experimental data on host factors affected by neurotropic flaviviruses were retrieved by literature mining and manually curated.

A table of curated studies with experimental evidence for flavivirus host interaction is displayed.

The columns are described below:

1. PMID – PubMed ID of the corresponding publication
2. Full text – full text document of the study
3. Virus – virus studied
4. Experimental method – experimental approach used in the study
5. Comment – manual curation comments
6. Host system – system used for the experimental measurement e.g. cell line
7. No of host factors identified – number of host genes experimentally associated with the virus
8. No of samples – number of samples analysed in the study
9. Validation of data quality – approach to quality assessment e.g. number of replicates
10. Platform – technical platform used for the measurements
11. Availability – availability of experimental data
12. Inclusion – Study inclusion criteria
13. 1st Author – first author of the study publication

14. Title – publication title
15. Journal – journal of publication
16. Year – publication year

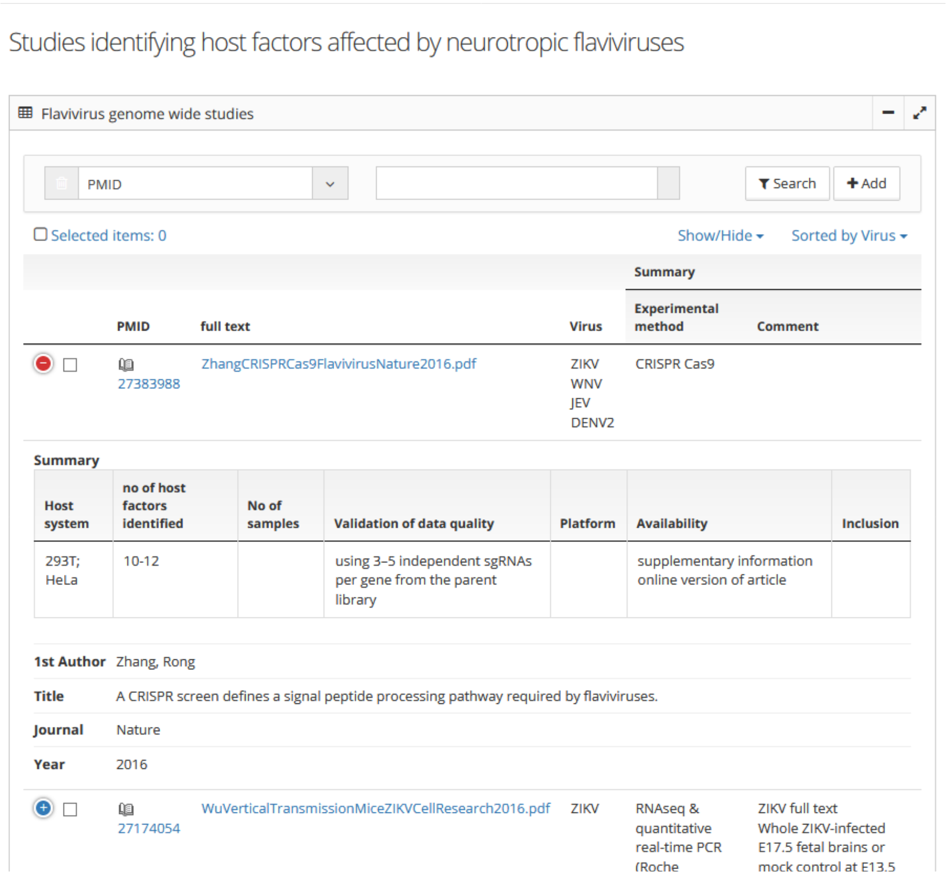

**Studies with flavivirus challenge** – The table displays detailed information about experimental studies to identify flavivirus interacting host factors. Note the red “-“ indicating expansion of columns initially hidden due to a small screen.

##### 2.3.8. Studies with virus challenge (neuro cells)

Select the **Zika KB** link in the top bar and then the green **Study with virus challenge (neuro)** button.

Studies providing experimental data on transcriptional effects of flavivirus challenges in neuronal cells which were retrieved by literature mining and manually curated.

The columns are described in 2.3.7.

#### 2.4. Disease maps

Select the **Zika KB** link in the top bar and then the **Disease maps** link in the left menu.

The **Disease maps** page provides access to expert curated network representations of all available molecular, cellular and phenotypic information about a biological process or disease in a structured, computationally accessible form. Biological concepts such as molecules, cells or phenotypes form the network nodes while their associations such as “activates”, “causes” for the edges between them.

Disease maps allow experts to collaborate online on the creation and refinement of mechanistic descriptions of biological processes. The maps available in the Zika KB are based on existing knowledge of molecular and cellular processes using literature mining as a foundation. In a second step protein-protein-interactions from publicly available databases, like BioGRID, have bene added. Finally all information used to populate the disease map can be reviewed and displayed as a network

Importantly, any kind of experimental or clinical data can then be superimposed on the created maps.

The available disease maps are displayed as a table. The table can be used as described in Section 1.4 Working with tables.

The columns are described below:

1. View Disease map – clicking on the disease map name will open the list of nodes and edges that describe the corresponding biological process. Clicking on the links within this column redirects to the disease map report and the subsequent network representation
2. No of disease map items – number of nodes involved in the biological process
3. No of disease map relations – number of edges involved in the biological process

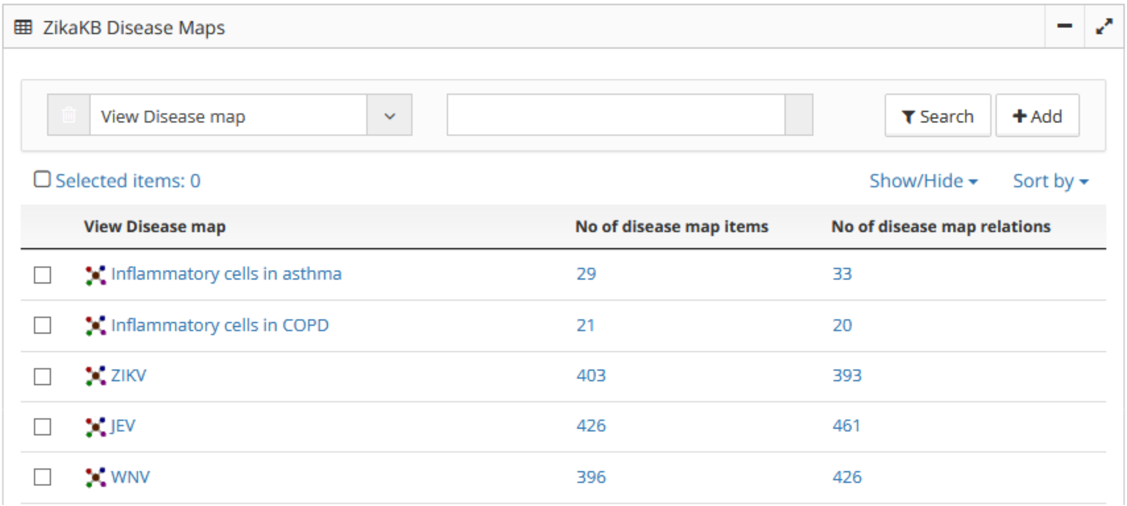
ZikaKB Disease Maps

View Disease map | Search | Add

Selected items: 0 | Show/Hide | Sort by

| View Disease map | No of disease map items | No of disease map relations |
|---|---|---|
| Inflammatory cells in asthma | 29 | 33 |
| Inflammatory cells in COPD | 21 | 20 |
| ZIKV | 403 | 393 |
| JEV | 426 | 461 |
| WNV | 396 | 426 |

**Disease maps** – The table displays the list of available disease maps with links to detailed information.

##### 2.4.1. Disease map report

Provides a detailed report of the selected disease map, listing all nodes and edges involved.

The results are displayed as a table with two separate sections for nodes and edges. A graphical or network display is available by using the “Graph” action menu on top of the page.

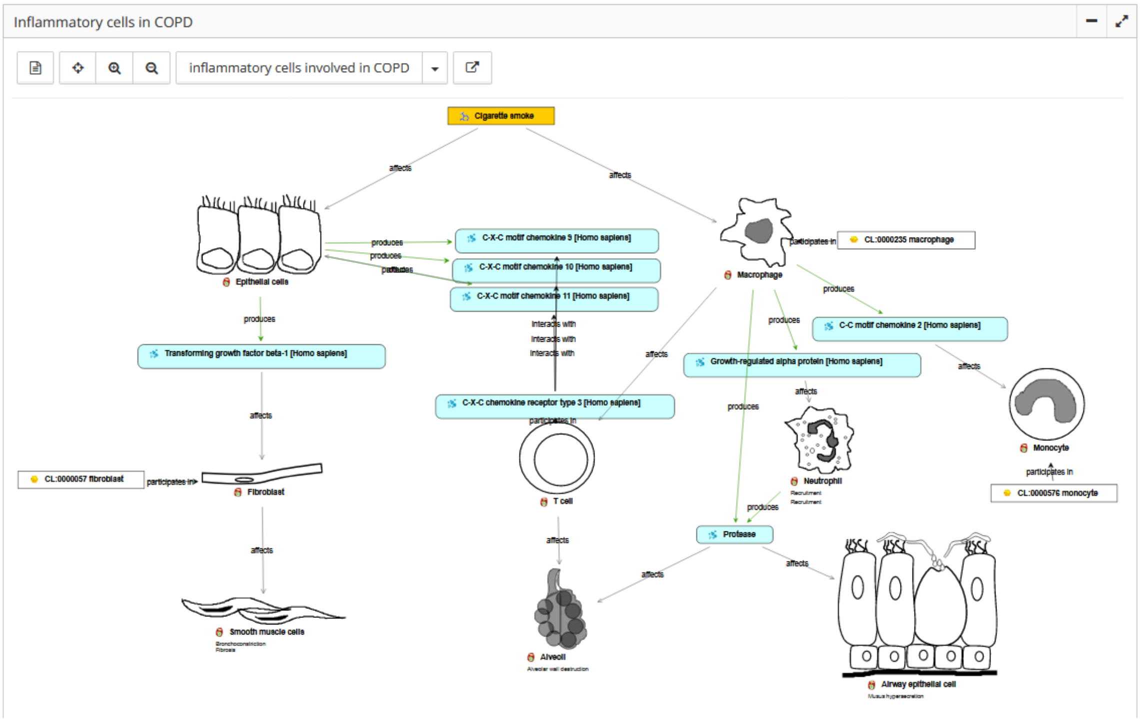

**Disease map graph** – The graph visualizes the nodes and edges involved in the described biological process.

The node columns are described below:

1. Object — the name of the node
2. Object type – the biological category of the node e.g. protein, cell type

The edge columns are described below:

1. Belief — the association of two nodes extracted from literature or structured databases provided as subject – predicate – object statement
2. Subject - the node forming the subject of the association
3. Object – the node forming the object of the association
4. Analysis – if the association was retrieved by literature mining rather than from expert knowledge or structured databases then the literature mining analysis from which the association was derived
5. Predicate – the predicate that was originally recognized by literature mining
6. Sentence – the sentence from which the association was originally extracted by literature mining
7. Lit Source – the publication the association was originally extracted from by literature mining
8. PMID – the PubMed ID of the publication
9. Ranking – the number of different publications supporting the association

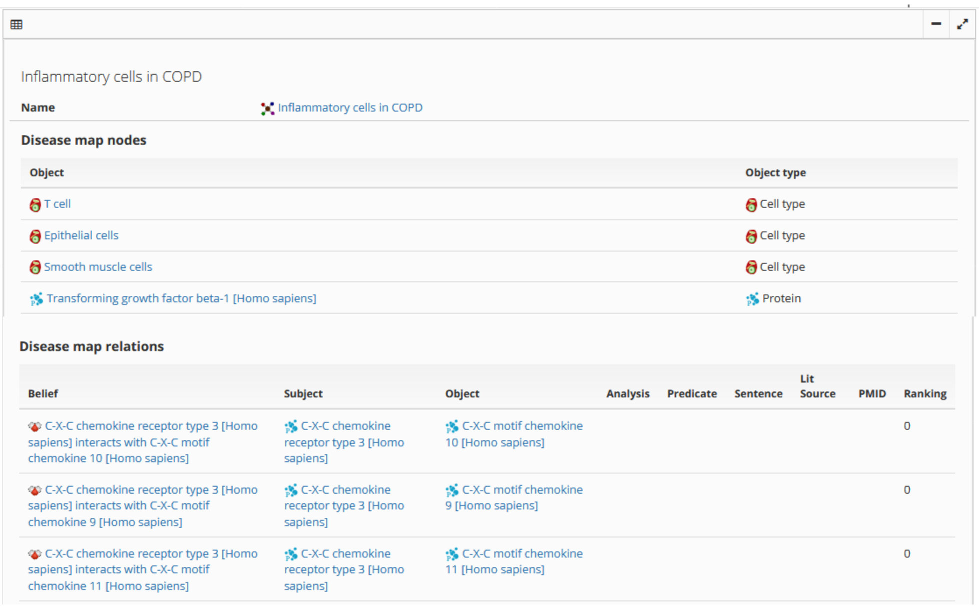

**Disease map report** – The table displays the two sections of the disease map report which separately list the map nodes and edges.

#### 2.5. VirHostNet Interactions

The **VirHostNet Interactions** page integrates a number of virus - host molecular interactions into the Zika KB that are provided by the virhostnet database.

##### 2.5.1. VirHostNet Interactions

Select the **Zika KB** link in the top bar and then the **VirHostNet Interactions** link in the left menu**.**

Lists all virus – host molecular interactions available from VirHostNet.

The results are displayed as a table. The table can be used as described in Section 1.4 Working with tables.

The columns are described below:

1. Source — the source molecule of the interaction
2. Target — the target molecule of the interaction
3. Source Gene — the gene encoding the source interacting protein
4. Process Source – the biological processes associated to the source molecule
5. Target Gene — the gene encoding the target interacting protein
6. Process Target – the biological processes associated to the target molecule
7. No of exp evidences – number of independent publications reporting the interaction
8. Taxon Interactor Source – the organism producing with the source molecule
9. Taxon Interactor Target – the organism producing with the target molecule

10. Interaction type – molecular interaction ontology derived interaction type
11. Method – molecular interaction ontology derived experimental method to identify the association
12. DB source – the structured database providing the interaction information
13. PMID – PubMed ID of the interaction reference

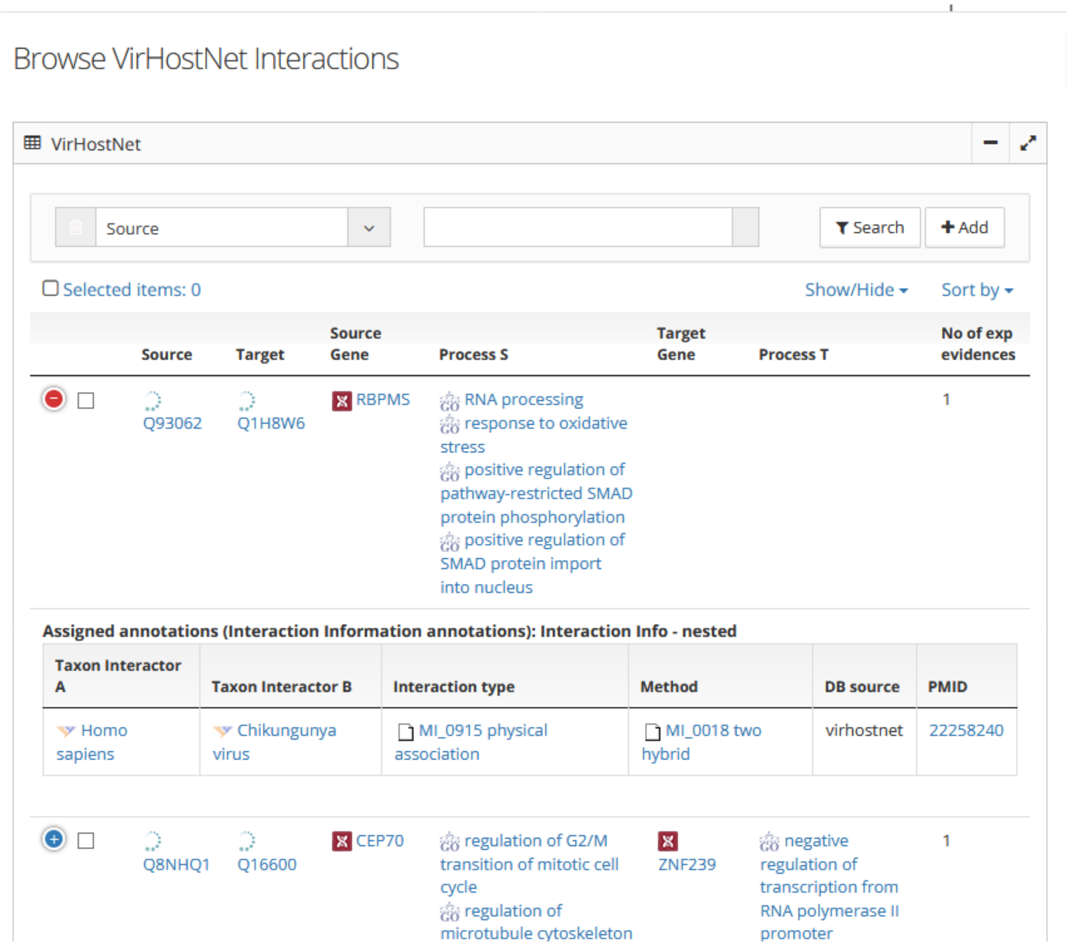

**VirusHostNet Interaction** – The table displays the virus – host molecular interactions integrated into the Zika KB from VirHostNet DB.

##### 2.5.2. Search for Host-Virus interactions by organism

Select the **Zika KB** link in the top bar and then the **VirHostNet Interactions** link in the left menu**.**

Search all virus – host molecular interactions available from VirHostNet for specific viruses and/or hosts.

To perform your own search select the button **Search for Host-Virus interactions by organism**:

1. Select the organism of interest from the "Organism" search field drop-down list

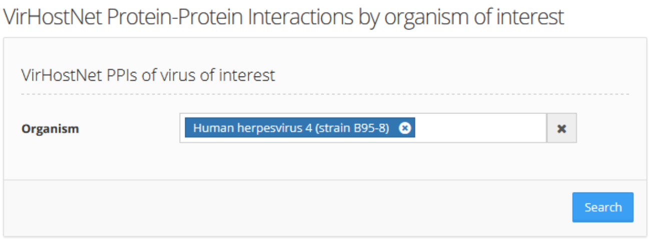

The results are displayed as a table. The table can be used as described in Section 1.4 Working with tables.

The columns are described in section 2.5.1.

##### 2.5.3. Browse Flavivirus VirHostNet Interactions

Select the **Zika KB** link in the top bar and then the **VirHostNet Interactions** link in the left menu**.**

Browse all virus – host molecular interactions available from VirHostNet pre-filtered for Flaviviruses.

Click the **Browse Flavivirus VirHostNet interactions** button on the main page to display a table of molecular virus – host interactions.

The results are displayed as a table. The table can be used as described in Section 1.4 Working with tables.

The columns are described in section 2.5.1.

#### 2.6. Host virus Interactions

The **Host Virus Interactions** page provides access to manually curated Host-Virus interactions with published experimental evidence which are accessible by individual buttons and are described in detail below.

##### 2.6.1. Browse Host-Virus Interactions

Select the **Zika KB** link in the top bar and then the **Host Virus Interactions** link in the left menu**.**

Lists host – virus protein-protein interactions integrated from multiple general protein-protein interaction databases into the Zika KB.

Click the **Browse Host-Virus Interactions** button on the main page to display the list of host – virus protein-protein interactions.

The results are displayed as a table. The table can be used as described in Section 1.4 Working with tables.

The columns are described below:

1. Source Gene — the gene encoding the source interacting protein
2. Source Protein — the source protein of the interaction
3. Target Gene — the gene encoding the target interacting protein
4. Taxon Interactor A – the organism producing with the source molecule
5. Taxon Interactor B – the organism producing with the target molecule

6. Interaction type – molecular interaction ontology derived interaction type
7. Method – molecular interaction ontology derived experimental method to identify the association
8. DB source – the structured database providing the interaction information
9. PMID – PubMed ID of the interaction reference
10. No of experimental evidence – the number of independent publications indicating the protein-protein interaction

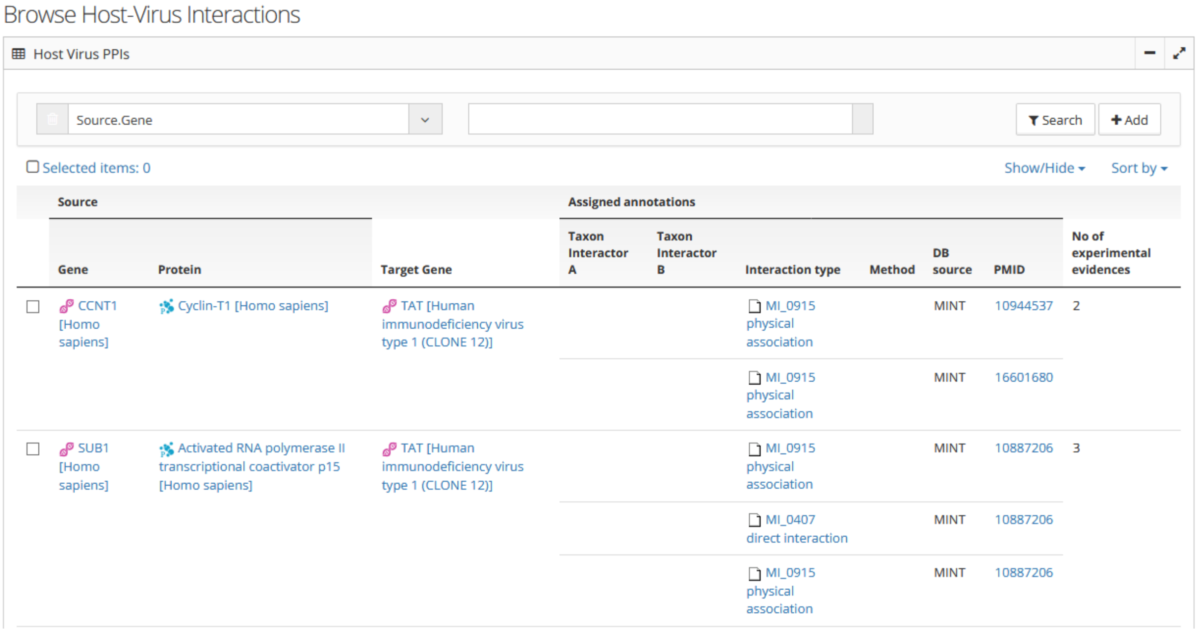

**Browse Host Virus Interactions** – The table displays the host-virus protein-protein interactions integrated from a set of protein-protein databases into the Zika KB.

##### 2.6.2. Search for Host-Virus Interactions by Virus

Search all integrated host – virus protein-protein interactions for a specific virus.

Click the **Search for Host-Virus Interactions by Virus** button on the main page to open a search form.

To perform your own search:

1. Select the virus(es) of interest from the “Virus” search field drop-down list

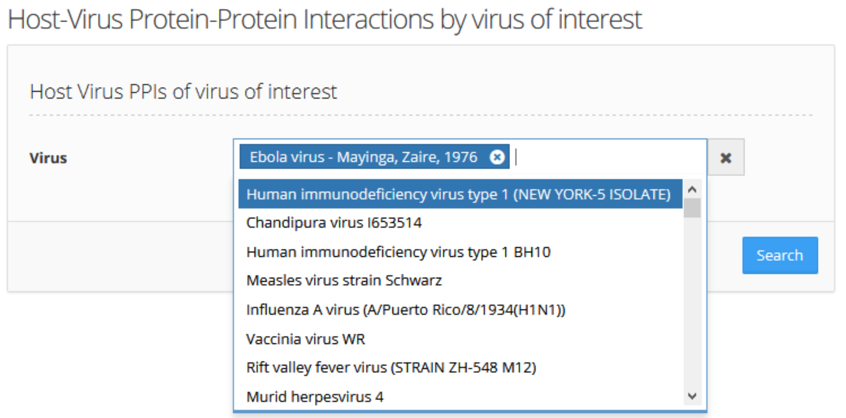

The results are displayed as a table. The table can be used as described in Section 1.4 Working with tables.

The columns are described in section 2.6.1.

##### 2.6.3. Search for Host-Virus Interactions by Virus (union/NOT)

Search all integrated host – virus protein-protein interactions for interactions occurring with a specific set of viruses but NOT others, for example to distinguish virus strains based on differential interactions.

Click the **Search for Host-Virus Interactions by Virus (union/NOT)** button on the main page to open a search form.

To perform your own search:

1. Select the virus(es) of interest from the “Virus” search field drop-down list
2. To exclude certain virus(es) select them in the “and NOT in Virus” search field drop-down list

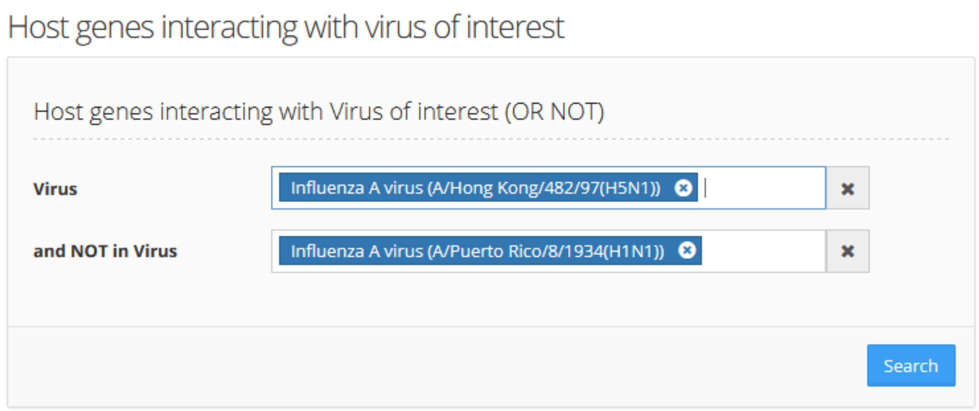

The results are displayed as a table. The table can be used as described in Section 1.4 Working with tables.

The columns are described in section 2.6.1.

##### 2.6.4. Search for Host-Virus Interactions by Virus (intersection)

Search all integrated host – virus protein-protein interactions for interactions occurring with a specific set of viruses but NOT others, for example to distinguish virus strains based on differential interactions.

Click the **Search for Host-Virus Interactions by Virus (union/NOT)** button on the main page to open a search form.

To perform your own search:

3. Select the virus(es) of interest from the “Virus” search field drop-down list
4. To exclude certain virus(es) select them in the “and NOT in Virus” search field drop-down list

The results are displayed as a table. The table can be used as described in Section 1.4 Working with tables.

The columns are described in section 2.6.1.

#### 2.7. Literature mining

Select the **Zika KB** link in the top bar and then the **Text Mining** link in the left menu**.**

The **Literature mining** page provides several tools for exploring the literature via literature mining.

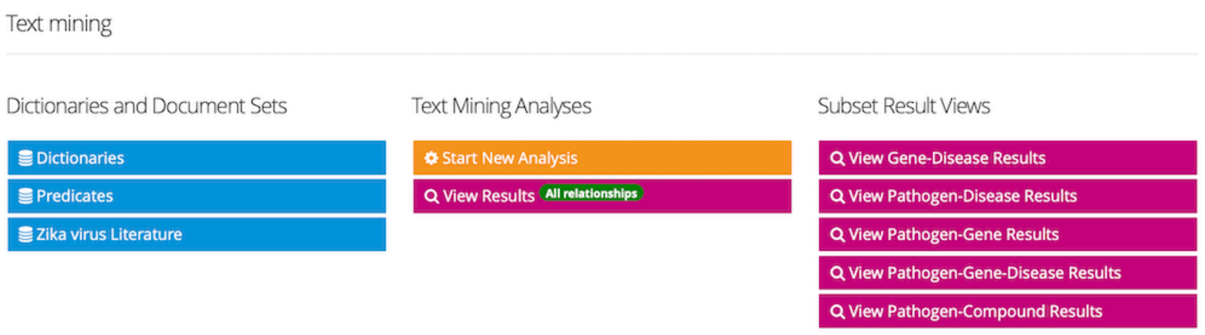

**Literature mining page**

The following buttons are available:

- Dictionaries — Types of vocabularies based on public ontologies (compounds OK
- Predicates — Types of predicates available for literature mining analysis
- Zika virus literature— View the list of Medline abstracts currently associated with Zika
- **Start Text Mining** — Start a new literature mining analysis
- **View Results** — View all literature mining results
- **View Gene-Disease Results** — View a subset of text-mining results restricted to gene-disease associations
- **View Pathogen-Disease Results** — View a subset of text-mining results restricted to pathogen-disease associations

- **View Pathogen-Gene Results** — View a subset of text-mining results restricted to pathogen-gene associations
- **View Pathogen-Gene-Disease Results** — View a subset of text-mining results restricted to pathogen-gene-disease associations
- **View Pathogen-Compound Results** — View a subset of text-mining results restricted to pathogen-compound associations
-
-

##### 2.7.1. Starting a new literature mining analysis

To start a new literature mining analysis, click the **Start New Analysis** button and complete the form:

1. Select the document corpus/repository on which the literature mining should be performed (option to select from: Medline abstracts, content from databases such as OMIM, ClinicalTrials. content from WHO news feeds, or subset of Medline abstracts based on a defined query), e.g. BioRS entries::MEDLINE
2. Select one or more dictionaries that are to be included in the mining, e.g. “Human Genes”, “Organism”, “Disease NR” (non redundant)
3. Enter a search key to restrict your search or **‘*’** for no restriction. Enter whole words or use wildcards ('*') to abbreviate words (keyword filters for content including entered keyword of interest). This will filter the set of MEDLINE abstracts for the entered search term, e.g. “Zika”
4. In the “Text Mining Quality” drop-down menu, select the type of search to be performed: syntax or a sentence-based.
    - **syntax** — for the best results in terms of precision
    - **sentence** — for a larger number of results, but less precision
5. Optionally, check the “Document Search” option. By default, relations and objects are only matched if they are in the same sentence. Activating this option widens the scope to the entire (!) document. This produces a larger number of results, but is less precise.
6. Optionally, check the “No Synonyms” option. By default, all synonyms of the entered input search term will be used to find matching documents. Activating this option narrows the result to only the documents that contain the entered search term.
7. Optionally, check the “Refresh Dictionaries” option. If the dictionaries are modified in any way, this option should be activated to make the changes available to the literature mining analysis.
8. Optionally, check the “Document Number Warning” option. Activating this option prompts the system to show warnings regarding the number of documents.
9. Click the “Run” button at the top of the form to start the analysis.
10. The task can be run in the background by clicking the “Run in background” button within the modal window

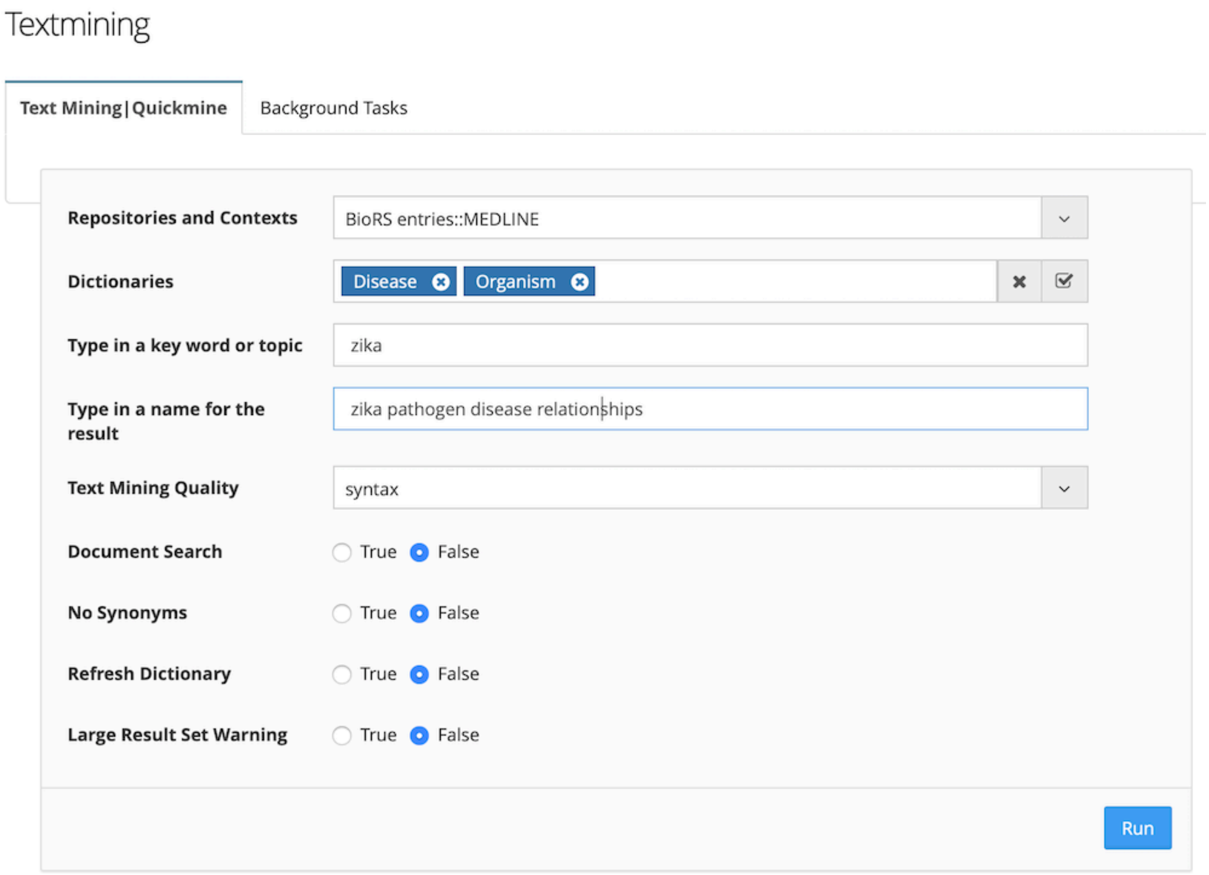

**Example of a literature mining analysis**

##### 2.7.2. Displaying literature mining results

To display a table of the literature mining analysis results, click the **View Results** button in the **Text Mining** page.

To view your result of interest:

1. Select the literature mining of interest from the “choose literature mining results” search field drop-down list (by default the “ZIKV curated 2018” set is selected, click the “select all” icon next to the cross to view all available literature mining analyses, the last result is the most recent one)
2. Click the **Search** button to view your selected analysis.

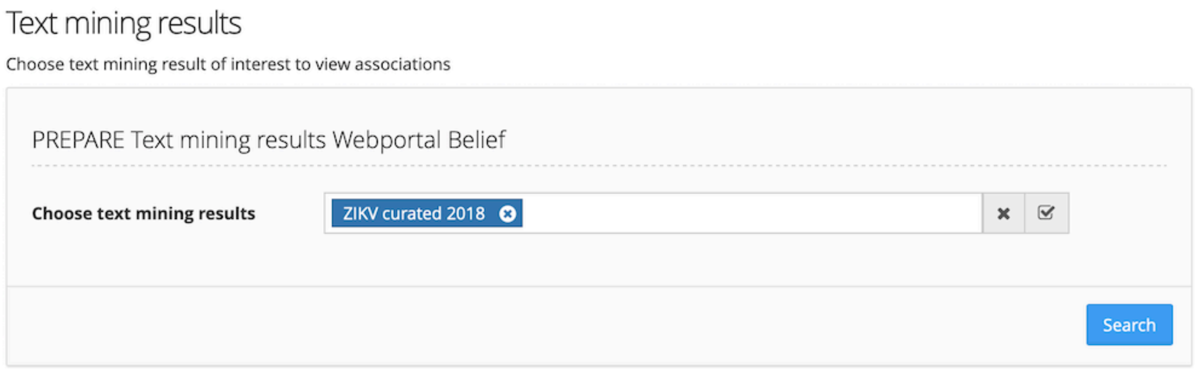

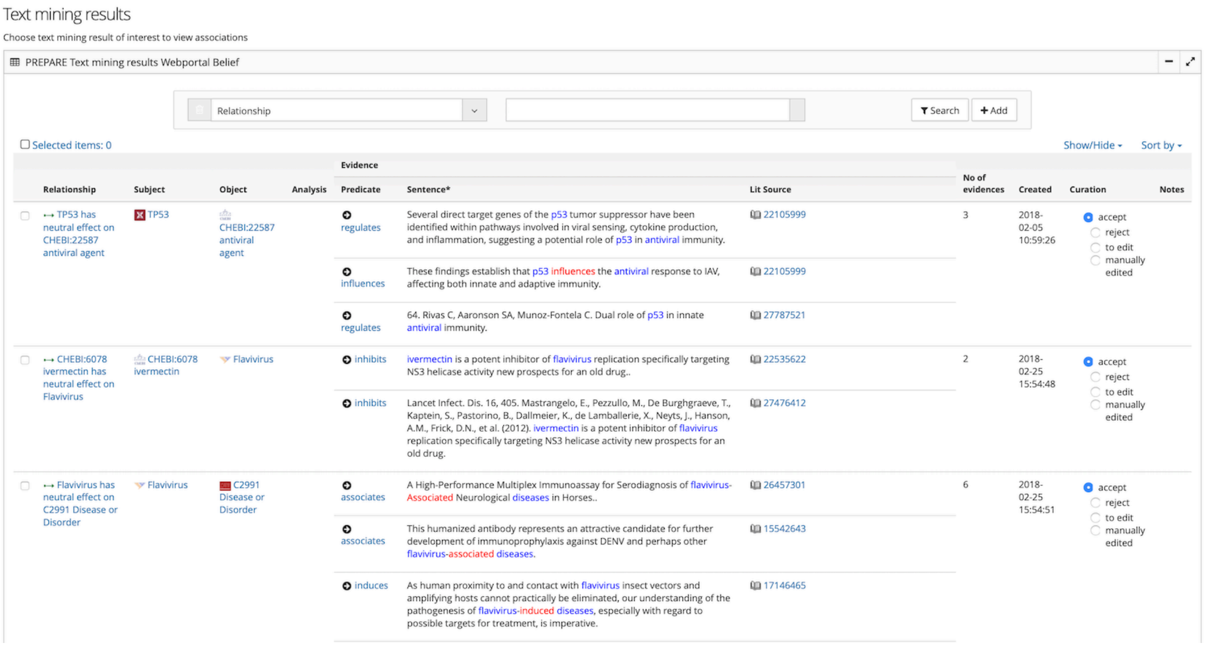

| | | | | Evidence | | | | | | |
|---|---|---|---|---|---|---|---|---|---|---|
| Relationship | Subject | Object | Analysis | Predicate | Sentence* | Lit Source | No of evidences | Created | Curation | Notes |
| TP53 has neutral effect on CHEBI:22587 antiviral agent | TP53 | CHEBI:22587 antiviral agent | | regulates | Several direct target genes of the p53 tumor suppressor have been identified within pathways involved in viral sensing, cytokine production, and inflammation, suggesting a potential role of p53 in antiviral immunity. | 22105999 | 3 | 2018-02-05 10:59:26 | accept / reject / to edit / manually edited | |
| | | | | influences | These findings establish that p53 influences the antiviral response to IAV, affecting both innate and adaptive immunity. | 22105999 | | | | |
| | | | | regulates | 64. Rivas C, Aaronson SA, Munoz-Fontela C. Dual role of p53 in innate antiviral immunity. | 27787521 | | | | |
| CHEBI:6078 ivermectin has neutral effect on Flavivirus | CHEBI:6078 ivermectin | Flavivirus | | inhibits | ivermectin is a potent inhibitor of flavivirus replication specifically targeting NS3 helicase activity new prospects for an old drug.. | 22535622 | 2 | 2018-02-25 15:54:48 | accept / reject / to edit / manually edited | |
| | | | | inhibits | Lancet Infect. Dis. 16, 405. Mastrangelo, E., Pezzullo, M., De Burghgraeve, T., Kaptein, S., Pastorino, B., Dallmeier, K., de Lamballerie, X., Neyts, J., Hanson, A.M., Frick, D.N., et al. (2012). ivermectin is a potent inhibitor of flavivirus replication specifically targeting NS3 helicase activity new prospects for an old drug. | 27476412 | | | | |
| Flavivirus has neutral effect on C2991 Disease or Disorder | Flavivirus | C2991 Disease or Disorder | | associates | A High-Performance Multiplex Immunoassay for Serodiagnosis of flavivirus-Associated Neurological diseases in Horses.. | 26457301 | 6 | 2018-02-25 15:54:51 | accept / reject / to edit / manually edited | |
| | | | | associates | This humanized antibody represents an attractive candidate for further development of immunoprophylaxis against DENV and perhaps other flavivirus-associated diseases. | 15542643 | | | | |
| | | | | induces | As human proximity to and contact with flavivirus insect vectors and amplifying hosts cannot practically be eliminated, our understanding of the pathogenesis of flavivirus-induced diseases, especially with regard to possible targets for treatment, is imperative. | 17146465 | | | | |

The results are displayed as a table. The table can be used as described in Section 1.4 Working with tables.

The columns are described below:

1. Relationship — the extracted triple consisting of a “subject”, “predicate” and “object”
2. Subject — the “subject” of the extracted triple
3. Object — the “object” of the extracted triple
4. Predicate — type of association retrieved by literature mining e.g. “regulates”, “influences”, “inhibits”
5. Sentence — sentence the triple was extracted from
6. Lit Source — PubMed ID or full text enry from which the relationship was retrieved
7. No of evidences — number of literature sources the relationship was extracted from
8. Created — creation time of the extracted relationship
9. Curation — curation status of et relationship
10. Notes — any notes curator has added

##### Displaying literature mining results

The literature mining results can be searched as follows:

1. Select the column in which to search from the drop-down menu above the table.
2. Enter the search term in the field next to the "Search" button.
3. Click the "+" (plus) button to add additional search terms and columns.

4. Click the "Search" button.
5. The search results will be displayed in the table.

##### Displaying Dictionaries and Predicates

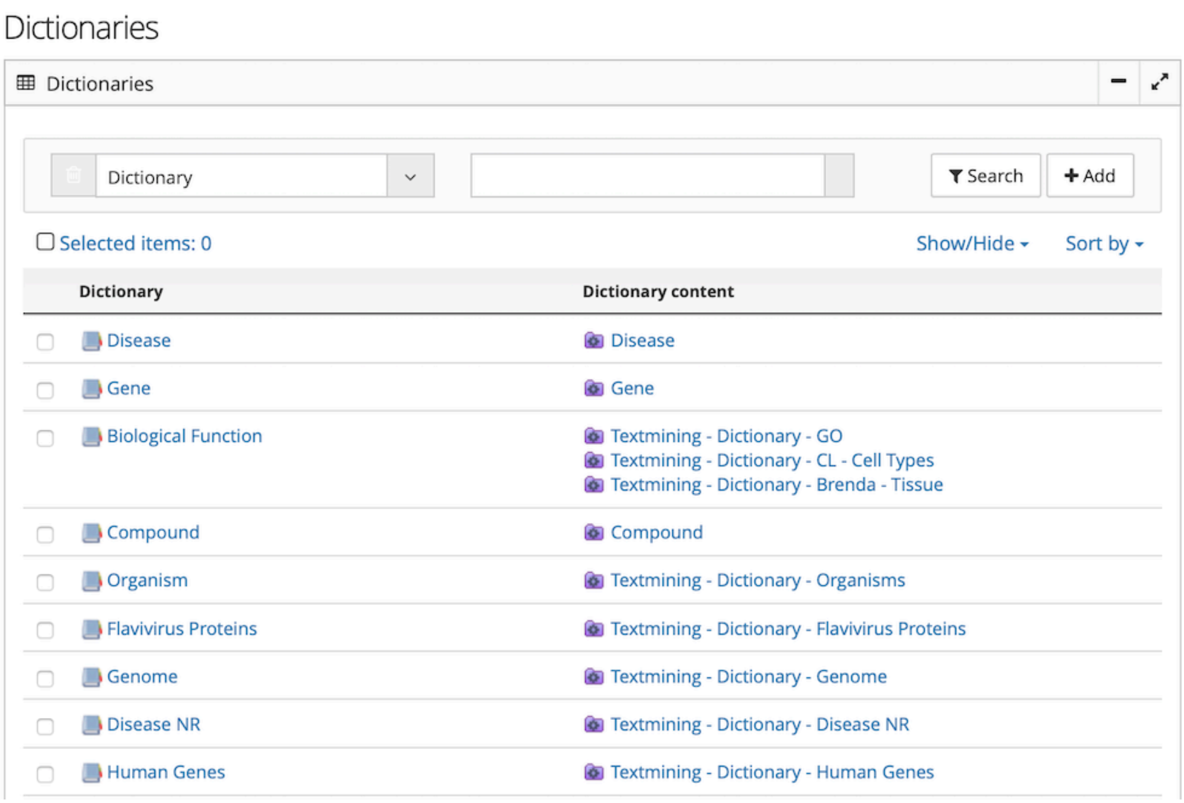

Types of vocabularies based on public ontologies

Predicates

Portal - Predicates - List

Predicates | Search | Add

Selected items: 0 | Show/Hide | Sort by

| | Predicates | Synonyms | No of relationships |
|---|---|---|---|
| | targeting | targeting | 0 |
| | treating | treating | 0 |
| | predicate | predicate | 0 |
| | hallmarking | hallmarking | 0 |
| | indicating | indicating | 0 |
| | not predicts | not predicts | 0 |
| | | | 0 |
| | is | is | 5505 |
| | surrounds | surrounds | 0 |
| | touches | touches | 0 |
| | starts | starts | 10 |
| | secretes | secretes<br>secrete<br>secreted<br>secreting<br>secretion | 0 |

Types of predicates available for literature mining analysis
