## Supplementary figures and images for "Informing epidemic (research) responses in a timely fashion by knowledge management - a Zika virus use case"

### Supplemental Figure 1

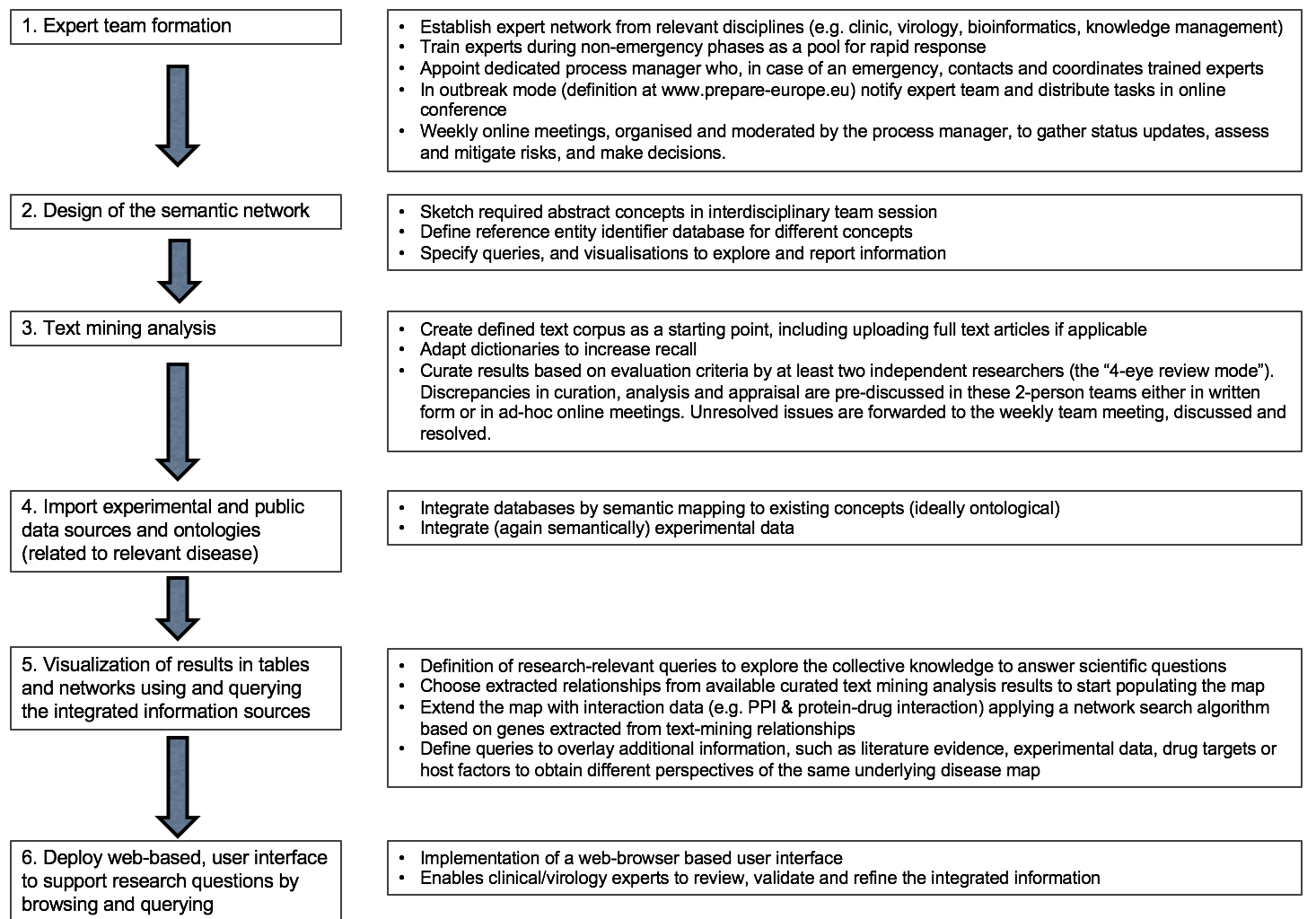

### Supplemental Figure 2

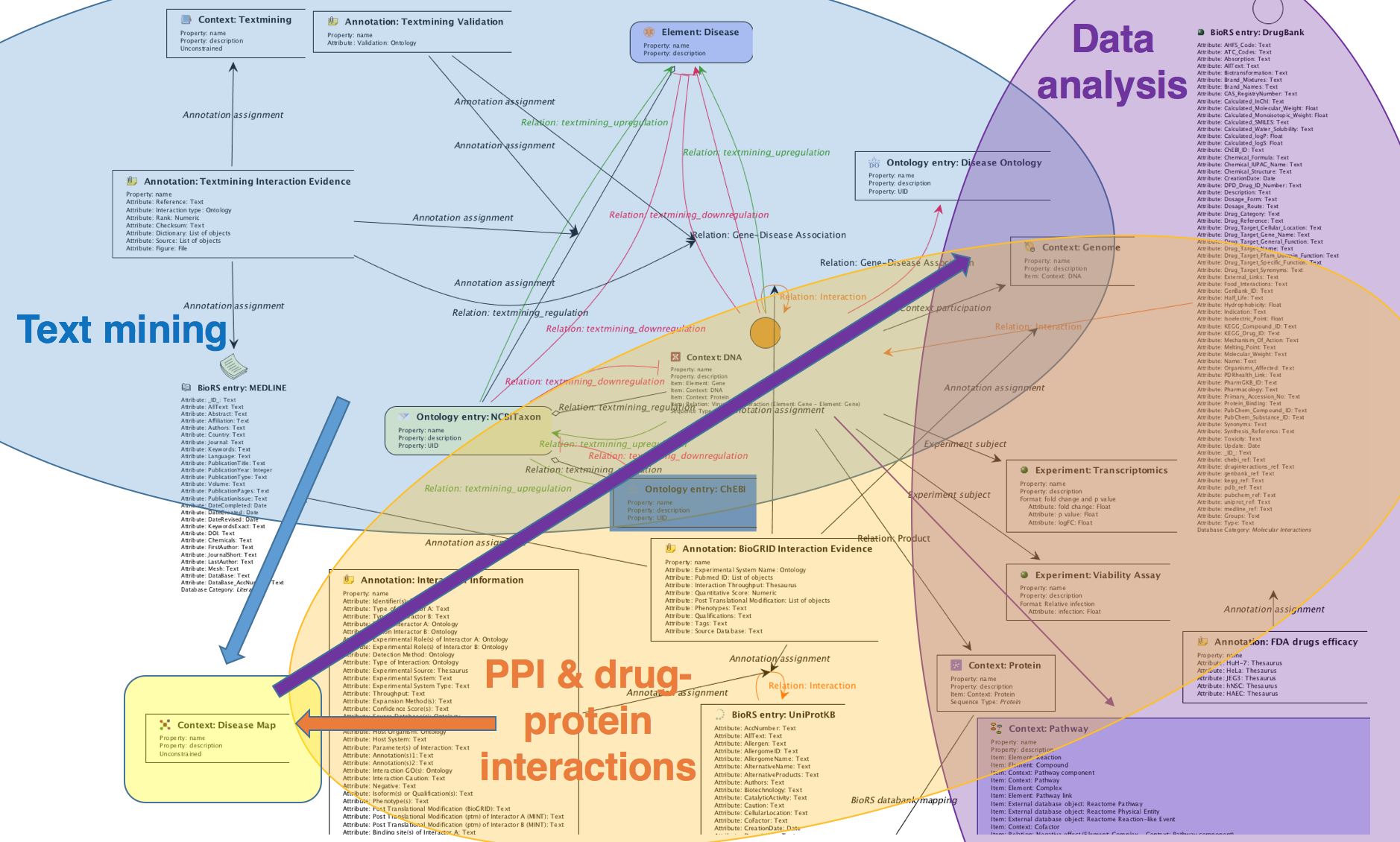

### Supplemental Figure 3

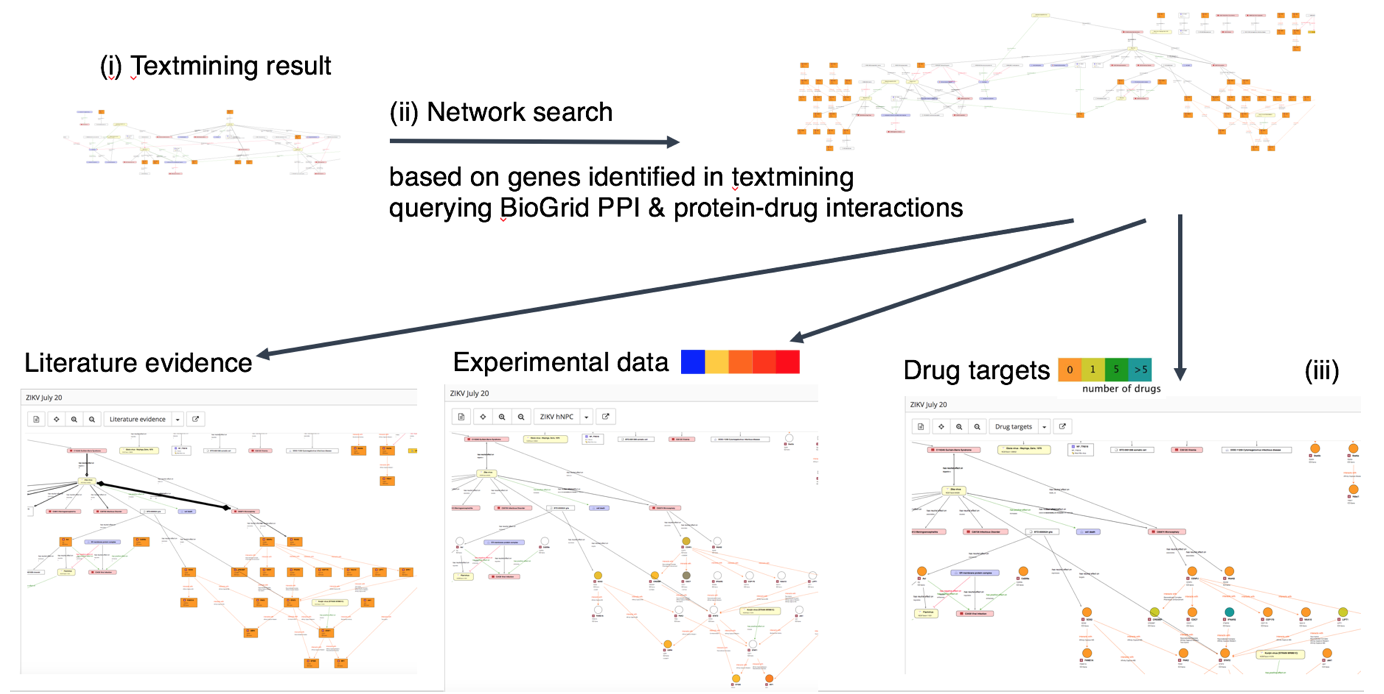
